# Apical PAR-3 caps orient the mitotic spindle in *C. elegans* early embryos

**DOI:** 10.1101/2023.03.27.534341

**Authors:** Naomi J. Stolpner, Nadia I. Manzi, Thomas Su, Daniel J. Dickinson

**Affiliations:** Department of Molecular Biosciences The University of Texas at Austin 2415 Speedway, PAT 206 Austin, TX 78712

## Abstract

During embryonic development, oriented cell divisions are important for patterned tissue growth and cell fate specification. Cell division orientation is controlled in part by asymmetrically localized polarity proteins, which establish functional domains of the cell membrane and interact with microtubule regulators to position the mitotic spindle. For example, in the 8-cell mouse embryo, apical polarity proteins form caps on the outside, contact-free surface of the embryo that position the mitotic spindle to execute asymmetric cell division. A similar radial or “inside-outside” polarity is established at an early stage in many other animal embryos, but in most cases it remains unclear how inside-outside polarity is established and how it influences downstream cell behaviors. Here, we explore inside-outside polarity in *C. elegans* somatic blastomeres using spatiotemporally controlled protein degradation and live embryo imaging. We show that PAR polarity proteins, which form apical caps at the center of the contact free membrane, localize dynamically during the cell cycle and contribute to spindle orientation and proper cell positioning. Surprisingly, apical PAR-3 can form polarity caps independently of actomyosin flows and the small GTPase CDC-42, and can regulate spindle orientation in cooperation with the key polarity kinase aPKC. Together, our results reveal a role for apical polarity caps in regulating spindle orientation in symmetrically dividing cells and provide novel insights into how these structures are formed.

## Introduction

During development, oriented cell divisions are important to ensure that cells are correctly positioned and adopt the proper fates during tissue specification. To carry out oriented cell divisions, positioning of the mitotic spindle must be coupled to cell polarity, which is the asymmetric distribution of cellular components that provides directionality to a cell. Cell polarity proteins localize to distinct domains on the plasma membrane, where they can orient both asymmetric and symmetric cell divisions in diverse developmental contexts, such as *Drosophila* sensory organ progenitors and neuroblasts, mouse skin and neuroepithelial progenitors, and *C. elegans* zygotes^1, 2^. In oriented cell divisions, polarity proteins in these domains interact with spindle orientation machinery (especially NuMA and the dynein-dynactin complex) to specify a division orientation through polarized force generation on the spindle. Distinct groups of polarity proteins can exclude one another from the cortex via biochemical interactions, thereby creating opposing polarity domains which ensure that polarized cell behavior occurs in specific areas of the cell^1, 3^.

One well-characterized example of oriented cell division in early animal embryos is inside-outside polarity in 8-cell mouse embryos^4^. Here, distinct groups of polarity proteins localize on the outside, cell-cell contact free (apical) membrane and the inside, cell-cell contact (basolateral) membrane. These proteins interact with spindle orientation machinery to localize one spindle pole toward the apical domain, resulting in an asymmetric cell division that specifies daughter cells of different fates^4–7^.

A similar inside-outside polarity is seen among many animal early embryos once they reach a multicellular stage^8–14^. In the *C. elegans* embryo, somatic blastomeres develop inside-outside polarity beginning at the 8-cell stage of development^9^. In 26-cell gastrulating embryos, inside-outside polarity is important to specify the apical surface of the endodermal precursors in order for apical constriction and cell internalization to occur^9, 15, 16^. However, the purpose of inside-outside polarity has not been explored in earlier 8-cell embryos, when it first emerges. At the 8-cell stage, AB lineage somatic blastomeres are the first to polarize on an inside-outside axis. These blastomeres divide symmetrically and differentiate later^17^, and their position within the embryo is important for their lineage specification^18–21^. Whether and how inside-outside polarity regulates the development of these blastomeres before gastrulation is not understood.

While a developmental role for inside-outside polarity in pre-gastrula stages in *C. elegans* has not been identified, studies of PAR polarity protein regulation in post-zygotic embryos have elucidated regulatory mechanisms for inside-outside polarity^9, 15, 22–24^. From this work, we know that the zygotic arrangement of anterior and posterior PAR proteins rearranges, with the anterior PARs (aPKC, PAR-3, PAR-6) localized to the outside, contact-free membrane and the posterior PARs (PAR-1, PAR-2) localized to cell-cell contacts^25, 26^. The scaffold protein PAR-3 can localize independently of at least some other PARs at these stages, and is necessary for aPKC/PAR-6 localization^25^. At cell contacts, the RhoGAP PAC-1 promotes apical localization of PAR proteins via a mechanism that was proposed to involve the small GTPase CDC-42^27, 28^. Together, these mechanisms ensure that gastrulating cells localize PARs apically to regulate apical constriction; loss of PAR-3 at the apical/outside surface results in gastrulation defects^16, 25^. How these polarity mechanisms might regulate developmental behaviors other than gastrulation in earlier stages is an interesting area of investigation.

Here, we set out to investigate the function and regulation of inside-outside polarity in the earliest *C. elegans* cells that polarize in this pattern, the 8-cell stage AB blastomeres. Using lineage-specific degradation and live imaging of endogenously tagged proteins, we found that PAR-3 and aPKC regulate oriented symmetric cell divisions in AB blastomeres at the 8-cell stage, when inside-outside polarity first appears. The aPARs localize to the outside surface in apical caps that are dynamic during the cell cycle and localize closely with centrosomes during spindle orientation. PAR-3 directs this behavior and is able to localize apically independently of myosin flows, microtubules, and other PAR proteins. The key polarity kinase aPKC localizes apically in a manner that is regulated by both PAR-3 and PAC-1, but is independent of CDC-42. Together, we propose a new role for inside-outside polarity in pre-gastrulation *C. elegans* embryos, and identify separable functions for PAR-3 in spindle orientation and in assembly of the apical PAR complex.

## Results

### PAR-3 forms apical caps that are closely associated with centrosomes

To study how PAR proteins establish inside-outside polarity, we focused on the AB lineage of cells, because they are the first to polarize inside-outside, starting at the 8-cell stage; progress through the cell cycle in synchrony; and are straightforward to image. In the zygote, PAR polarity is dynamic over the cell cycle ^29–34^, so we began by performing time-lapse imaging of endogenously tagged mNeonGreen::PAR-3 in the four ABxx blastomeres from their birth to their division. During mounting, embryos can randomly orient with either their left or right side towards the coverslip, so that typically only two (left side) or three (right side) AB blastomeres are visible at a time (Figure 1A). We imaged embryos in both orientations for every experiment, and observed equivalent behaviors in all four AB cells except where noted below. As AB cells completed the 4-to-8 cell division, PAR-3 localized evenly on the cortex with no visible polarity. As the four AB cells progressed through the cell cycle, PAR-3 became enriched apically, on the exterior surface of the cells, and was excluded from cell-cell contacts, in agreement with previous work^25, 26, 28, 35, 36^. Interestingly, PAR-3 did not localize uniformly over the apical surface, but rather formed distinct ‘caps’ in the center of each AB cell’s apical membrane (Figures 1B-C, Movie 1). These PAR-3 caps reached peak intensity approximately 5 minutes after ABxx birth, persisted until nuclear envelope breakdown (NEBD), and then dissipated as the cells progressed through mitosis. Apical caps formed and dissolved with similar timing and appearance in all four AB blastomeres. PAR-3 was visibly punctate in apical caps, suggesting that PAR-3 forms oligomers on the cortex at the 8-cell stage, as it does at the 1-cell stage^32, 37^.

**Figure 1:**
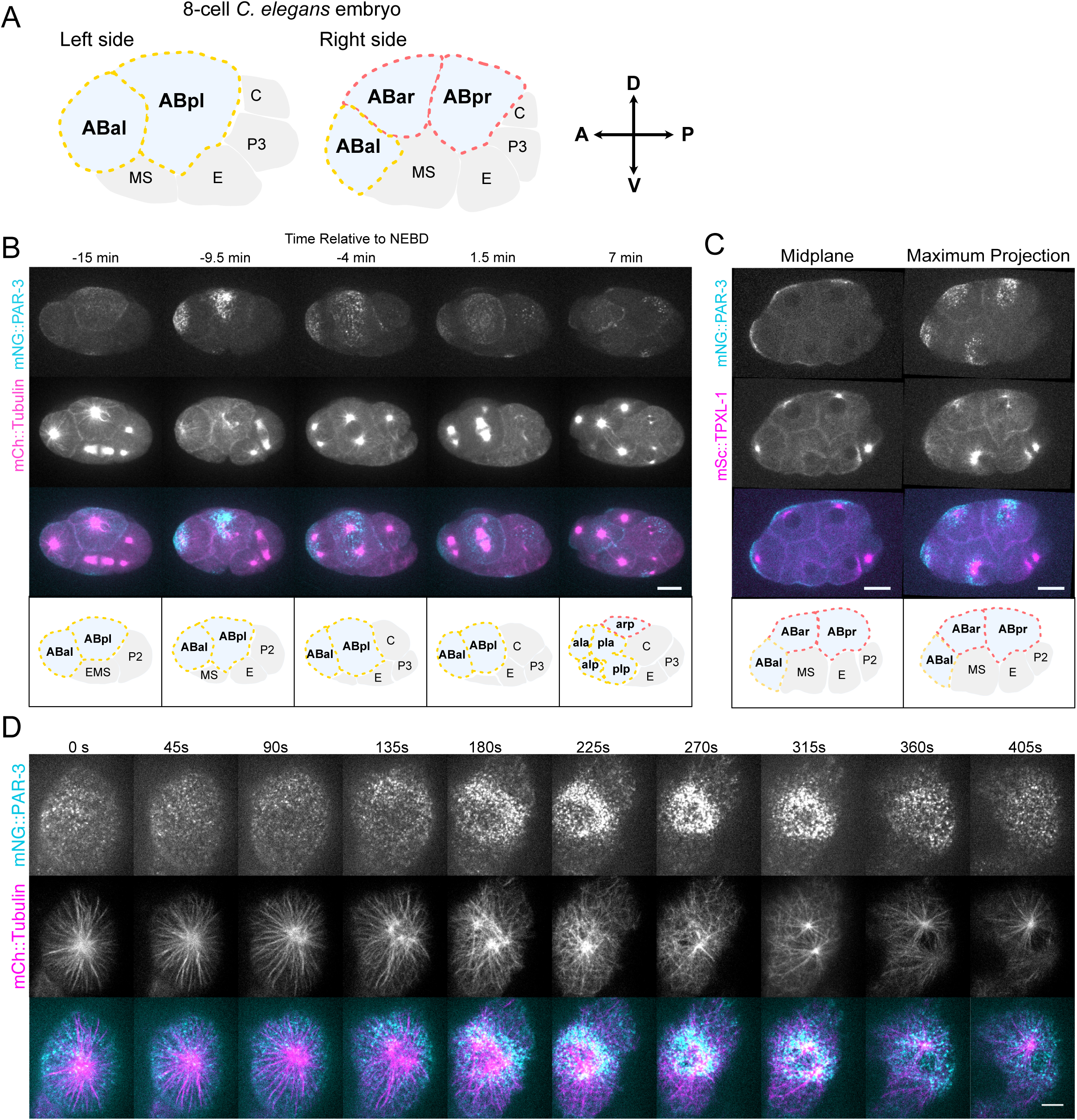
PAR-3 apical caps localize near centrosomes and are dynamic during the cell cycle. A. Cartoon of 8-cell embryos with left and right sides shown. Left AB blastomeres are outlined with yellow dashed lines and Right AB blastomeres are outlined in red dashed lines. Arrows indicate the Anterior/Posterior (A/P) and Dorsal/Ventral (D/V) axes, all laterally mounted embryos will be displayed with anterior to the left and dorsal at the top. Note that the ABal cell is visible from both sides due its position within the embryo. B. Time-lapse montage of embryo expressing mNG::PAR-3 and mCh::Tubulin from left side, max projections starting just after the ABxx cells are born. This embryo was imaged from the left side; the top two blastomeres, with mNG::PAR-3 signal at t = –9.5 min, are the ABxl blastomeres. Time is indicated relative to Nuclear Envelope Breakdown (NEBD). Scale bar represents 10 µm. C. Midplane slices & maximum projections of mNG::PAR-3 and TPXL::Sc (centrosome marker). This embryo was viewed from the right side; ABal, ABar and ABpr are visible as top-left three cells mNG::PAR-3 signal. Scale bar represents 10 µm. D. Time-lapse montage of embryo expressing PAR-3::mNG and mCh::Tubulin, mounted en-face dorsally with the ABal blastomere in view. Montage starts just after ABxx cells are born, displayed every 45 seconds. Scale bar represents 5 µm. See also: Movies 1-2.

We also imaged Tubulin::mCherry and the centrosomal marker mScarlet-I::TPXL-1 in these experiments, because centrosomes and microtubules in these cells undergo well-described rearrangements that provide a readout of cell cycle progression^38^. Unexpectedly, these images revealed a close association between PAR-3 caps and centrosomes (Figures 1B-C). As PAR-3 caps formed, newly duplicated centrosomes were visible near the apical surface, adjacent to the PAR-3 cap. PAR-3 domains expanded as centrosomes migrated to opposite sides of the nucleus to set up the next mitotic spindle (tubulin, t=3), and PAR-3 remained diffuse but still polarized over the mitotic spindle as the cells progressed into metaphase (t=4). Apical PAR-3 puncta largely dissipated by telophase/cytokinesis, when the AB cells divided symmetrically (t=5). We did not observe any asymmetric inheritance of PAR-3 caps or puncta.

To better visualize PAR-3 dynamics during apical cap formation and contemporaneous centrosome and microtubule movements, we imaged embryos that were mounted with the apical surface of AB cells (i.e., the dorsal side of the embryo) toward the coverslip (Figure 1D, Movie 2). At t=0, the 4-to-8 cell mitotic spindle was still in view, and PAR-3 clusters were evenly dispersed throughout the apical surface. PAR-3 began concentrating at the apical cap as the centrosome underwent fragmentation and duplication (t=90-270s) ^38^. PAR-3 caps appeared most concentrated as the centrosomes began migration to align the next mitotic spindle (t=315), and the PAR-3 domain expanded and moved across the cortex in tandem with the migrating centrosomes (t=360). Together, this data pointed to a possible physical connection between apical caps and centrosomes/MTs and led us to hypothesize that there may be a relationship between apical caps and the mitotic spindle.

### Loss of PAR-3 leads to spindle orientation defects

To examine a possible connection between apical PAR-3 caps and mitotic spindle orientation in the AB blastomeres, we used the ZF1 system^15, 39^ to degrade PAR-3 in the AB cell lineage. The ZF1 degron sequence, derived from the PIE-1 protein, is recognized by the E3 ubiquitin ligase ZIF-1 and targeted for degradation in somatic blastomeres, allowing depletion of PAR proteins in the AB lineage without interfering with PAR polarity the zygote stage. We examined a strain in which ZF1 was integrated into the *par-3* gene using Cas9-triggered homologous recombination^40, 41^, resulting in degradation of endogenous PAR-3 in the AB blastomeres beginning during the 4-cell stage (Figure 2B). We first looked for defects in cell division orientation in explants, generated by removing the eggshell at the 2-cell stage and isolating the AB cell, in order to exclude possibly conflicting spindle orientation cues from neighboring P-lineage cells or from confinement within the eggshell^19, 42^. We denote the developmental stage of these explants by indicating the number of blastomeres present followed by the total number of cells that would be present in an intact embryo; for example, 4_AB_/8 indicates a cluster of 4 AB cells that are the same age as those from an 8-cell embryo (Figure 2A). We examined the 4_AB_/8 to 8_AB_/12 cell division in explants, which corresponds to the first cell division after PAR-3 caps form.

**Figure 2:**
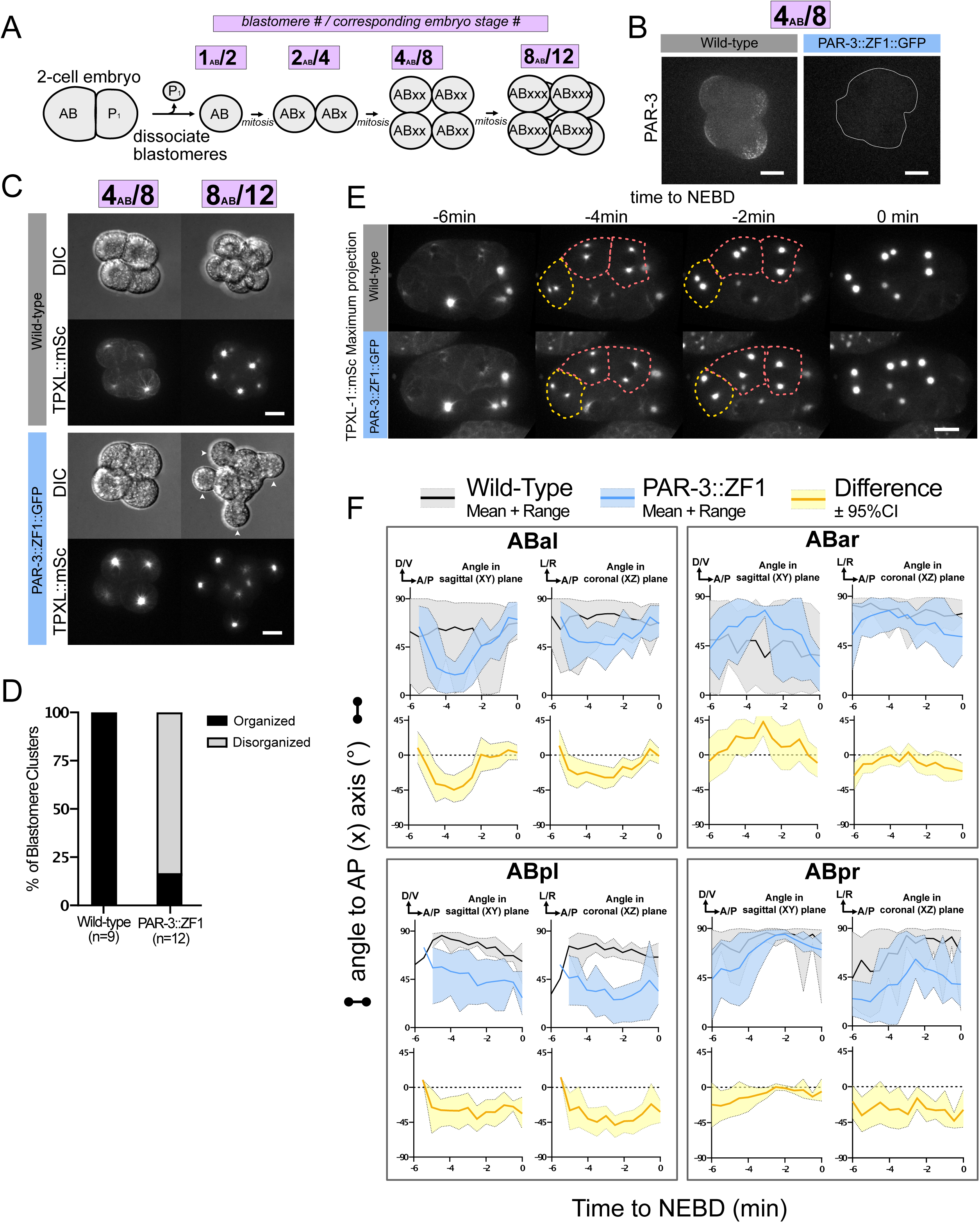
Loss of PAR-3 causes spindle orientation defect. A. Cartoon workflow of blastomere explant experiment. 2-cell embryos were dissociated to separate the AB cell from P_1_. AB was allowed to divide into ABx, then mounted and imaged through the next two cell divisions. Explants are named by the AB blastomere number in the cluster and which corresponding embryo stage they are equivalent to, displayed in purple boxes above each step. B. Representative images of 4_AB_/8 blastomere explants from wild-type mNG::PAR-3 embryo and PAR-3::ZF1::GFP embryo. In PAR-3::ZF1::GFP, blastomere explant is outlined in white. Scale bars represent 10 µm. C. Images from time-lapse live movies of blastomere explants from wild-type (mNG::PAR-3; TPXL-1::mSc) and PAR-3::ZF1::GFP (PAR-3::ZF1::GFP; TPXL-1::mSc) imaged from 4_AB_/8 to 8_AB_/12 stage. DIC single plane images and TPXL-1::mSc maximum projections representative of the majority of the dataset are shown. Arrowheads point to cells without neighbors. Scale bars represent 10 µm. D. Phenotype scoring of 8_AB_/12 blastomere explants from Fig2C. Dataset was blinded and scored based on whether the blastomere explant was organized/spherical or disorganized/had cells protruding from the cluster. E. TPXL-1::mSc maximum projection images from time-lapse live full-volume imaging of wild-type (mNG::PAR-3; TPXL-1::mSc) and PAR-3::ZF1::GFP (PAR-3::ZF1::GFP; TPXL-1::mSc) embryos. Right side is displayed, three ABxx cells are visible along the top and left of the embryo. The cells are outlined (colors match Figure 1) at times when centrosome positioning defects were observed. Time is shown relative to nuclear envelope breakdown (NEBD). Scale bars represent 10 µm. F. Quantification of spindle angle over time in AB blastomeres. Full-volume images were displayed in 3D, and centrosomes were annotated manually. X, Y, and Z coordinates were used to determine the XY and XZ angles of the spindle relative to the embryo’s AP axis over time, starting from the beginning of centrosome separation until NEBD. Time is shown relative to NEBD. See also: Figure S1, Movies 4-6.

Both wild-type and PAR-3::ZF1::GFP explants formed adherent clusters at the 4_AB_/8 cell stage (Figure 2B-C, Movies 3 and 4). Wild-type explants had PAR-3 caps on the contact-free surface that formed at the correct developmental time, while PAR-3::ZF1::GFP blastomeres had no detectable PAR-3::ZF1::GFP signal, indicating successful degradation of PAR-3 (Figure 2B). As the cells divided from the 4_AB_/8 to 8_AB_/12 cell stage, we observed a striking difference between wild-type and PAR-3::ZF1::GFP explants. In wild-type, mitotic spindles oriented parallel to one another and perpendicular to the inside/outside axis of the cluster, thereby maintaining a compact cluster shape through to the 8_AB_/12 stage (n=9/9) (Figure 2C, Movie 3), as reported previously for AB cell explants^20, 43, 44^. In PAR-3::ZF1::GFP explants, the spindles oriented radially along the inside-outside axis of the cluster, resulting in cell divisions that placed daughter cells on the edge of the cluster with no neighbors. To quantify these phenotypes, we blindly scored blastomere clusters as either “compact” (cells form a generally spherical cluster) or “disorganized” (presence of cells on the edges of a cluster with no neighbors) (Figure 2D, Movie 4). The disorganized phenotype was observed in the majority of PAR-3::ZF1::GFP explants but was never seen in wild-type.

To compare these observations to spindle positioning in intact embryos, we imaged centrosome positions over time in wild-type or PAR-3-depleted 8-cell embryos. In wild-type embryos, AB cells followed a stereotyped spindle orientation and division pattern, as expected (Figure 1A). Following the preceding cell division, centrosomes localized near the outside of each blastomere, near the PAR-3 cap. Centrosomes migrated to establish spindle orientation for the next cell division, with one centrosome (usually the ventral one) migrating more while the other centrosome remained closer to the cortex. ABal, ABpl, and ABpr aligned their spindles parallel to the dorsal/ventral axis of the embryo, while ABar shifted its spindle towards the left/right axis, a behavior that is known to depend on an external Wnt signal from the neighboring C cell^21, 45, 46^. In all AB cells, the centrosomes migrated directly to their final locations and the spindles did not reorient before division, except in some ABal blastomeres which oriented their spindles D/V as normal, then rotated the spindle 180° end-over-end, which still resulted in D/V orientation prior to division.

Consistent with our results in explants, depletion of PAR-3 from intact embryos resulted in a spindle positioning defect. Centrosomes pairs in each AB cell were aligned along an axis that pointed towards the contact with the neighboring EMS cell, as though arranged along spokes of a wheel with the EMS cell in the center (Figure 2E, Movies 5 and 6).

To quantify these defects in intact embryos, we collected full-volume Z stacks of embryos, made 3D reconstructions, and measured the spindle angles in each AB blastomere over time in both sagittal (X-Y) and coronal (X-Z) planes. We observed spindle angle defects when PAR-3 was degraded, which we broadly categorized into two groups based on the range of spindle angles observed in each blastomere. First, ABal and ABar wild-type spindle angles had a large range, while PAR-3 depletion resulted in a spindle angle defect in which the spindle position was more stereotyped (Figure 2F, top). Second, ABpL and ABpR had more consistent wild-type spindle angles, and also showed spindle angle defects of varying severity when PAR-3 was depleted (Figure 2F, bottom). Unlike in explants, in intact embryos some of these defects were corrected prior to cell division, resulting in largely normal spindle orientation at NEBD (Figures 2E-F) and normal cell positioning at the 12-cell stage (data not shown). We also measured centrosome separation from the same movies and found that it was unaffected (Figure S1). Together, this shows that PAR-3 is involved in spindle orientation in inside-outside polarized AB cells, though there is some redundant mechanism that ultimately corrects the final spindle orientation and cell division axis.

### Apical cap formation is independent of microtubules and myosin flows

Having shown that PAR-3 caps form in the 8-cell embryo and function in the AB lineage to orient mitotic spindles, we wanted to know how these PAR-3 caps form at the apical surface. PAR-3 caps formed in a cell-cycle dependent manner above the nucleus as centrosomes were fragmenting and rebuilding. This proximity between PAR-3, centrosomes, and microtubules led us to hypothesize that microtubules emanating from centrosomes toward the apical cortex might provide a spatial cue that specifies where PAR-3 apical caps should form. To test this hypothesis, we permeabilized embryos, added nocodazole to disrupt microtubules, and asked if PAR-3 caps formed normally. By monitoring the mCh::Tubulin signal, we verified that microtubules were no longer visible 60-90 seconds after nocodazole addition. Therefore, we added nocodazole after the 4-to-8 cell division was complete but at least 90 seconds before apical cap formation began.

Control embryos incubated with DMSO had normal tubulin appearance and normal PAR-3 caps (Figure 3A). In stage-matched nocodazole-treated embryos, microtubules were not visible and neighboring cells (EMS) were arrested in the cell cycle. Still, apical PAR-3 caps were able to form with similar timing and positioning on the apical surface as in control embryos. Consistent with the disruption of microtubules, centrosomes in treated AB cells were displaced away from the apical surface and towards the center of the embryo (Figure 3A, asterisks). These data indicate that PAR-3 apical caps can form even when apical microtubules and apical centrosome localization are disrupted.

**Figure 3:**
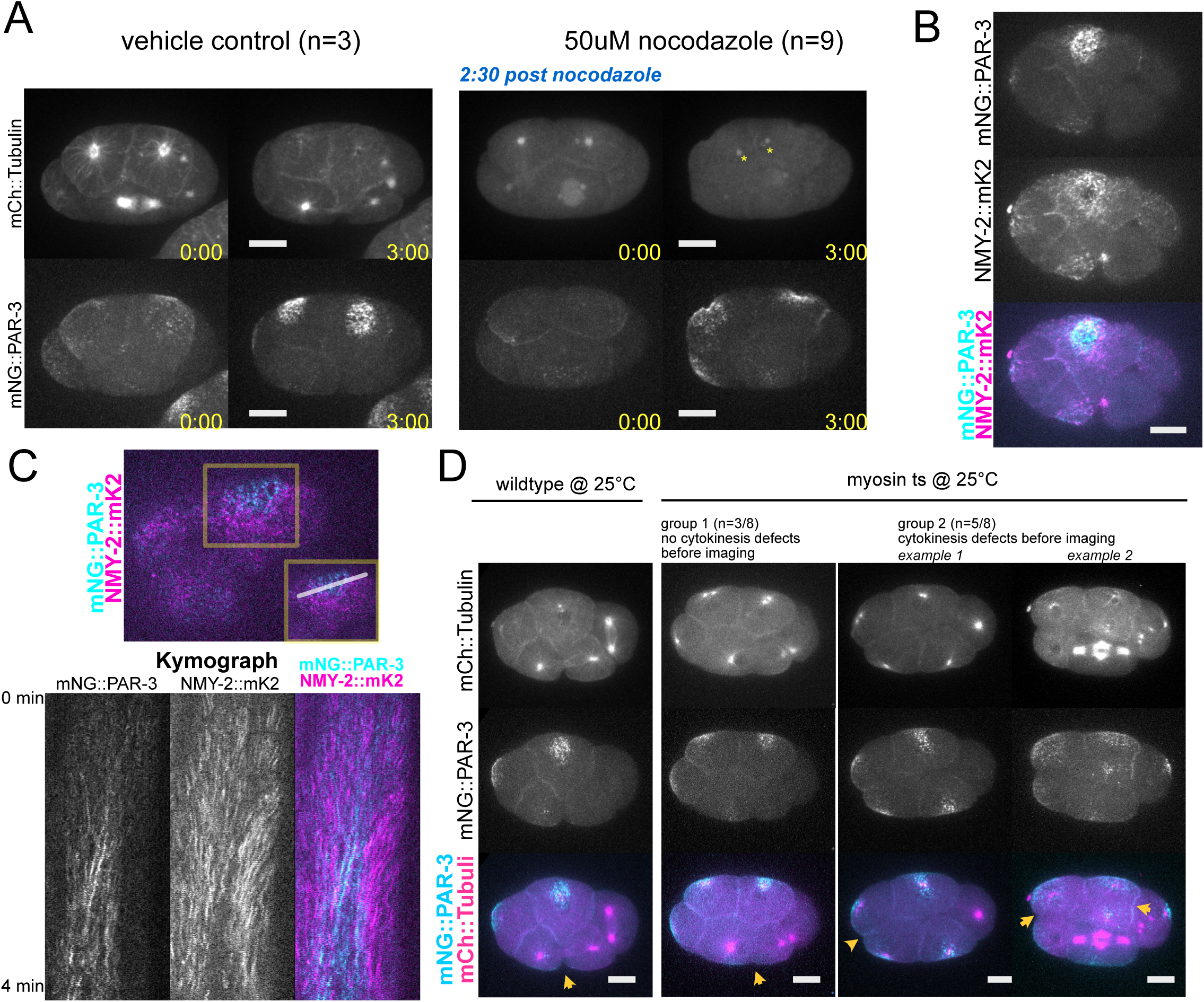
PAR-3 can polarize when myosin or microtubules are disrupted. A. Vehicle control and 50 µM nocodazole-treated embryos expressing mCh::Tubulin; mNG::PAR-3. embryos were permeabilized with *perm-1* RNAi and imaged in media + DMSO vehicle in an open coverslip device that allows drug addition during imaging. Right: Nocodazole treatment. After the 4-8 AB cell division, 50 µM nocodazole was added and wicked through the device; MTs were not visible 60-90 sec after addition. Left: For controls, no nocodazole was added. Right side of embryo is shown; ABxx blastomeres visible along top/anterior. Timepoints shown are equivalent staged embryos, 3 minutes elapsed between time points. Scale bars represent 10 µm. B. Maximum projection of embryo expressing NMY-2::mKate2; mNG::PAR-3 at peak apical cap formation. Left side of embryo is shown focused on Abal; ABxx blastomeres are visible along top/anterior. Scale bars represent 10 µm. C. Cortical image of embryo expressing NMY-2::mKate2; mNG::PAR-3 and kymograph. Image is at the beginning of mNG::PAR-3 apical cap formation. Inset shows selection across the ABpl apical cap used for kymograph. Kymograph displays pre-apical cap formation to peak apical cap intensity. Time is on Y axis. D. Maximum projection of control embryos (mCh::Tubulin; mNG::PAR-3) and myosin (ts) embryos (myosin (ts); mCh::Tubulin; mNG::PAR-3). Wild-type embryos formed normal cytokinesis furrows in neighboring mitotic cells (yellow arrow). myosin (ts) embryos had disrupted visibly shallow cytokinesis furrows and/or partial furrows that receded (yellow arrows). Embryos were divided into two groups. Group 1 (one example shown): embryos with cells that had normal centrosome numbers and therefore didn’t have a visible cytokinesis defect before imaging; Group 2 (two examples shown): embryos with cells that had duplicate centrosome numbers indicating a cytokinesis defect before imaging. Here, arrows indicate partial furrows that regressed or areas between two nuclei that had no membrane. Scale bars represent 10 µm. See also: Figure S2.

Because microtubule disruption did not block the formation of apical caps, we considered other cytoskeletal components that could contribute spatial cues for PAR-3 localization. The actomyosin cortex is critical for polarizing PAR-3 in the zygote via actomyosin flows^35, 37, 47^. Myosin has been shown to flow apically at the 8-cell stage^25, 35^ and we observed that myosin was enriched at the apical surface where it appeared to form a ring around the PAR-3 cap (Figure 3B). Thus, actomyosin flows appeared a likely candidate to transport PAR-3 to form apical caps in the 8-cell embryo.

First, to test whether PAR-3 and myosin flowed towards the apical membrane together as apical caps formed, we mounted embryos dorsally and imaged apical cap formation at the most cortical plane of embryos with tagged PAR-3 and myosin (NMY-2) (Figure 3C, top). From these timelapse movies, we made kymographs from a line across the widest part of the apical surface to visualize myosin and PAR-3 movement over time (Figure 3C, bottom). As previously reported, we observed myosin flows that moved away from cell-cell contacts and towards the center of the apical surface, appearing as sloped lines in the kymograph. The myosin signal started near cell-cell contacts at the edges of the apical surface and moved toward the center, ending in a ring with reduced myosin signal at the center. While PAR-3 clusters appeared on the cortex around the same time as myosin flows began, these PAR-3 clusters did not flow from the edges of the apical surface towards the center as myosin did (Figure 3C, bottom). Instead, the PAR-3 clusters formed near the center of the apical surface and persisted in the same location, with little evidence of flow/movement as seen by straighter lines on the kymograph. Together, these observations in wild-type embryos suggested that while myosin does flow apically, these flows are not obviously transporting PAR-3 clusters to form the apical cap.

We next wanted to directly test whether myosin flows were necessary for apical PAR-3 localization by perturbing myosin function imaging PAR-3 localization during apical cap formation. Because a null allele or RNAi knockdown of myosin would inhibit cell division in the zygote, halting development before the 8-cell stage, we used two parallel methods to perturb myosin while still allowing the embryo to develop to the 8-cell stage. First, we used a temperature sensitive myosin allele^48^. We dissected embryos at the permissive temperature of 15°C, then immediately imaged at the restrictive temperature 25°C. Embryos expressing mNG::PAR-3, mCh::Tubulin with wild-type myosin developed normally and formed apical PAR-3 caps after temperature shifting in this way (Figure 3D, left). We observed two classes of myosin ts embryos: Group 1 - embryos without cytokinesis defects before imaging (Figure 3D, column 2); and Group 2 - embryos that were multinucleate at start of imaging and produced extra centrosomes in the subsequent cell division, suggesting a prior cytokinesis defect (Figure 3D, columns 3 & 4). Group 2 embryos most likely result from embryos that failed to complete cytokinesis during the delay of several minutes between dissecting at 15°C and the start of imaging at 25°C.

In both groups of myosin ts embryos, we observed clear evidence of myosin disruption. Neighboring EMS blastomeres did not have defined cleavage furrows compared to wild-type (Figure 3D, bottom, yellow arrows), and AB cells had partial furrows that regressed (group 2, example 1, arrow) or were multinucleate with no membrane in between nuclei/centrosomes (group 2, example 2, arrow). In both groups, all cells failed to divide. Despite this myosin disruption, we saw apical PAR-3 caps that formed at the medial apical surface, positioned above nuclei and centrosomes. Multinucleate cells that had failed cytokinesis formed multiple caps, one adjacent to each nucleus and centrosome (Figure S2A).

We wanted to use a second parallel approach to perturb myosin, so we took advantage of the fact that different myosin regulatory kinases have distinct, semi-redundant roles and performed RNAi on *mrck-1*, a myosin activating kinase that is dispensable for zygotic polarity and cell division but is known to contribute to apical myosin enrichment in E lineage blastomeres at gastrulation^16, 37^. *mrck-1(RNAi)* embryos developed to the 8-cell stage and appeared grossly normal, but had disrupted apical myosin architecture at the 8-cell stage. At peak PAR-3 apical cap intensity in wild-type embryos, myosin appeared as a fine mesh with small puncta in the AB cells, whereas myosin in *mrck-1* RNAi treated embryos was diminished and appeared disorganized (Figure S2B-C). Still, in both wild-type and *mrck-1* RNAi embryos, PAR-3 caps formed at the medial apical surface. The total embryonic PAR-3 signal was lower in *mrck-1* RNAi embryos than wild-type (Figure S2B), which is most likely an RNAi artifact or a secondary consequence of MRCK-1 depletion that led to less mNG::PAR-3 expression. However, this reduction in PAR-3 levels did not affect the ability of PAR-3 to organize in an apical cap.

Together, these data argue against critical roles for microtubule-based trafficking or myosin flows in establishing apical PAR-3 caps, and suggest that PAR-3 polarization may be more dependent on interactions/relationships within the PAR complex itself. We therefore shifted our focus to understanding how PAR-3 interacts with other key polarity proteins to establish apical caps and orient mitotic spindles at the 8-cell stage.

### PAR-3 apical cap formation is upstream of aPKC

Our results thus far indicate that PAR-3 localizes independently of myosin and microtubules to form apical caps, where it plays a role in spindle orientation. We hypothesized that the polarity kinase aPKC, a PAR-3 binding partner, could be involved in PAR-3 cap formation and/or spindle orientation. In zygotic polarization, aPKC and PAR-3 have an interdependent relationship: PAR-3 polarizes only weakly and transiently when aPKC is depleted by RNAi, and aPKC does not polarize or localize to the membrane in the absence of PAR-3^49–51^. However, in several apicobasally polarized cell types, PAR-3 and aPKC can localize independently and/or have distinct functions^52–56^, so we wanted to directly test the relationship between aPKC and PAR-3 in apical caps.

In wild-type embryos, aPKC localized to apical caps (Figures 4A-B), in agreement with previous studies^9, 15, 22^. Interestingly, we also saw a significant pool of aPKC at cell-cell contacts that has not been previously noted (Figure 4A-B). We quantified aPKC localization by making line scan measurements from the apical cap throughout the cytoplasm to the contact. From these line scans, we found the apical and lateral (cell-cell contact) maximum pixel intensity values, and an average cytoplasmic value. These measurements confirmed that aPKC was enriched at cell-cell contacts compared to the cytoplasm (Figure 4C). Cell-cell contact localization of aPKC was much more prominent than that of PAR-3, which is barely detectable at contacts (Figures 1B-C and 4D-E), suggesting that there is a pool of aPKC at cell-cell contacts that is not associated with PAR-3. When PAR-3 was degraded, aPKC localized in the cytoplasm and did not display a polarized localization on any membranes (Figure 4B-C).

**Figure 4:**
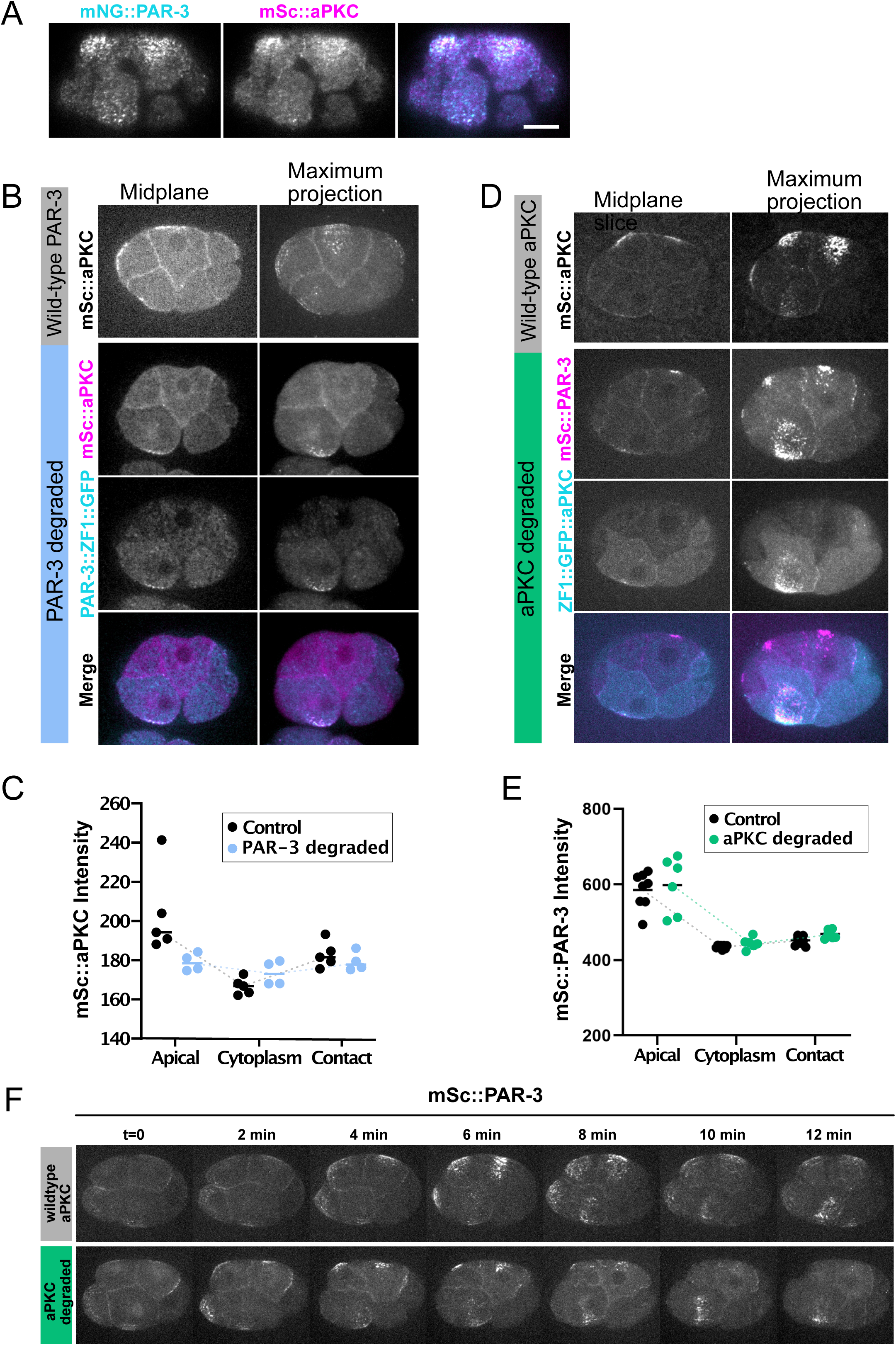
PAR-3 instructs aPKC localization and polarizes independently of aPKC during spindle orientation. A. Images of a wild-type embryo showing colocalization of aPKC and PAR-3 in apical caps. Scale bars represent 10 µm. B. Localization of Sc::aPKC and PAR-3::ZF1::GFP. Scale bars represent 10 µm. C. Quantification of data in B. Linescans were taken from midplane slices from the apical membrane through the cytoplasm and to the AB cell-cell contact. Apical maximum, cytoplasmic average (of 10 datapoints) and contact maximum were plotted. D. Localization of Sc::PAR-3 in aPKC::ZF1::GFP. Scale bars represent 10 µm. E. Quantification of data in D, as in C. F. Time-lapse of Sc::PAR-3 localization (max projection) in wildtype and aPKC::ZF1::GFP background. Scale bars represent 10 µm. See also: Figure S3

Next, we asked whether aPKC was required for apical localization of PAR-3. We imaged live embryos with endogenously tagged aPKC::ZF1::GFP and mSc::PAR-3 and quantified the localization using linescans from the cap to the contact, as we had done for aPKC localization. We found that when aPKC was degraded, PAR-3 still localized to apical caps (Figure 4D-E). Notably, PAR-3 was still polarized in the absence of aPKC and did not localize to cell-cell contacts. We observed punctate PAR-3, suggesting that PAR-3 oligomerization could still occur. However, apical PAR-3 caps were narrower in the absence of aPKC, which could indicate that aPKC might tune the amount of PAR-3 or contribute to maintenance of the PAR-3 cap. This agrees with previous observations of fixed, immunostained embryos showing that PAR-3 could still localize apically without aPKC at later developmental stages^15^. We also examined the dynamics of PAR-3 caps following aPKC depletion and observed that PAR-3 caps formed with similar timing, but dissolved slightly earlier in the cell cycle (Figure 4F). It is possible that this reflects a role for aPKC in maintenance of PAR-3 caps, or alternatively, caps may appear to dissolve sooner because they are smaller to begin with. Regardless, these data show that aPKC plays a measurable but not strictly essential role in supporting PAR-3 localization to apical caps.

Together, these data show that PAR-3 is upstream of aPKC at this stage of development. PAR-3 can still polarize to apical caps even when aPKC is degraded, suggesting that unlike in the zygote, the cues that polarize PAR-3 are aPKC-independent. On the other hand, aPKC localization to the plasma membrane at both caps and contacts requires PAR-3.

### aPKC loss also causes spindle orientation defects

We next asked whether aPKC localization downstream of PAR-3 is required for mitotic spindle orientation in the AB lineage. To test this, we examined spindle orientation after depletion of endogenous aPKC via the ZF1 system (Figure 5A). In 4_AB_/8 explants from ZF1::GFP::aPKC embryos, we observed disorganized cell clusters similar to PAR-3::ZF1, but with reduced frequency and severity. Disorganized clusters still represented a majority of samples, but typically had only 1-2 cells protruding from the cluster rather than 4 (Figure 5B-C, compare to 2C, Movie 7). In intact ZF1::GFP::aPKC embryos, we also saw similar defects as in PAR-3::ZF1, with spindles tilted toward the contact with the EMS cell in the center of the embryo (Figures 5D-E, compare to 2E-F, Movie 8).

**Figure 5:**
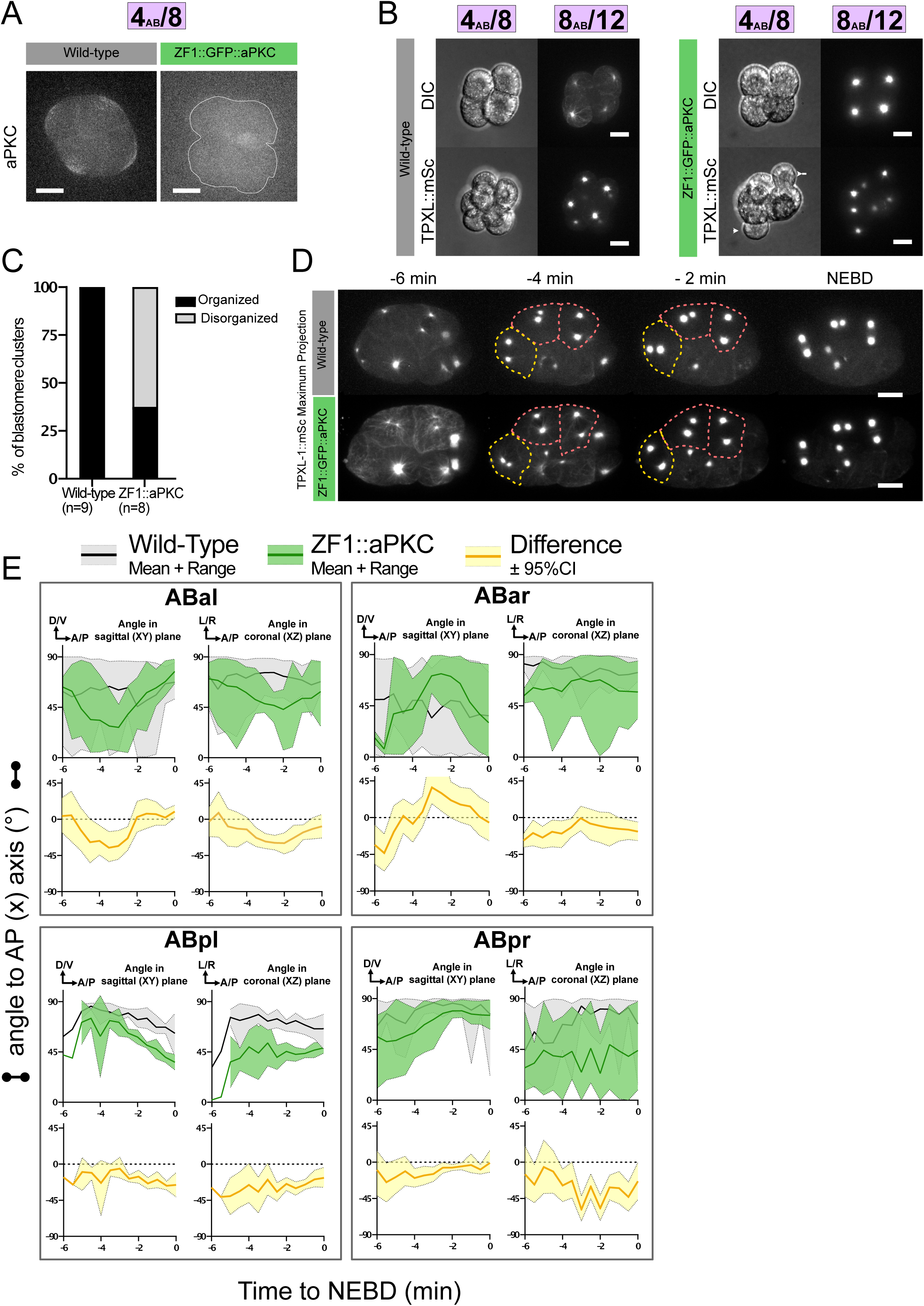
aPKC degradation results in a mild spindle orientation phenotype. A. aPKC localization in wild-type (mNG::aPKC; TPXL-1::mSc) and ZF1::GFP::aPKC (ZF1::GFP::aPKC; TPXL-1::mSc) 4_AB_/8 blastomere explants as in figure 2. In ZF1::GFP::aPKC, blastomere explant is outlined in white. Scale bars represent 10 µm. B. Representative images of 4_AB_/8 blastomere explants from wild-type mNG::aPKC embryo and ZF1::GFP::aPKC embryo. Wild-type data is the same as in Figure 2, shown again to facilitate comparison. Arrowheads indicate cells protruding from the cluster. Scale bars represent 10 µm. C. Phenotype scoring of 8_AB_/12 blastomere explants from Fig5B. Dataset was blinded and scored based on whether the blastomere explant was organized/spherical or disorganized/had cells protruding from the cluster. D. TPXL-1::mSc maximum projection images from time-lapse live full-volume imaging of wild-type (mNG::aPKC; TPXL-1::mSc) and ZF1::GFP::aPKC (ZF1::GFP::aPKC; TPXL-1::mSc) embryos. Right side is displayed, three ABxx cells are visible along the top and left of the embryo. ABpl spindle is visible in the center of the embryo starting at t=8 due to projection of full-volume image. Displayed is every 2 minutes, images taken every 30 seconds. The cells are outlined (colors match Figure 1) at times when centrosome positioning defects were observed. Scale bars represent 10 µm. A. E. Quantification of spindle angle in ABal from Fig5D. Full-volume images were displayed in 3D, and centrosomes were annotated manually. X, Y, and Z coordinates were used to determine the XY and XZ angles of the spindle relative to the embryo’s AP axis over time, starting from the beginning of centrosome separation until NEBD. Interval = 30sec. See also: Movies 7-8.

Together with the localization experiments in Figure 4, these results suggest two possible interpretations regarding the roles of PAR-3 and aPKC in spindle orientation.First, it is possible that PAR-3 controls AB spindle orientation in an aPKC-dependent manner. Alternatively, PAR-3 may act on spindles via an aPKC-independent mechanism, and the spindle phenotypes in ZF1::aPKC could be a secondary consequence of smaller apical PAR-3 caps in the absence of aPKC (Figure 4D). To distinguish between these possibilities, we decided to perturb other known regulators of aPKC localization, aiming to disrupt aPKC localization independently of PAR-3.

### Apical aPKC is regulated by PAC-1 independently of CDC-42

aPKC alone does not bind stably to the plasma membrane; it localizes via membrane binding proteins including PAR-3 and the small GTPase CDC-42. PAR-3 and CDC-42 form independent complexes with aPKC/PAR-6, which are believed to have distinct roles in polarity in the zygote^50, 51, 57, 58^. Previous work on *C. elegans* inside-outside polarity has emphasized a putative role for active, GTP-bound CDC-42 as a key regulator of aPKC localization^22–24^. This model is based on genetic identification of a RhoGAP protein, PAC-1, which localizes exclusively to cell-cell contacts and not the apical/outside surface^22^. As a putative CDC-42 GAP, PAC-1 is thought to promote CDC-42 GTP hydrolysis, resulting in inactive, GDP-bound CDC-42. Because PAC-1 localizes to cell-cell contacts, active CDC-42 was proposed to accumulate and then recruit aPKC/PAR-6 at the outside/apical cell membrane, which is free of PAC-1. Consistent with this model, loss of PAC-1 resulted in uniform localization of aPKC/PAR-6 at both cell-cell contacts and contact-free membranes^22^. We therefore decided to use PAC-1 depletion as an alternative, PAR-3-independent means to perturb aPKC localization at the 8-cell stage.

First, we used RNAi to knock down *pac-1,* and measured the amounts of aPKC at caps and cell-cell contacts. In *pac-1(RNAi)* embryos, aPKC was unpolarized, with equivalent levels at the apical membrane and at cell-cell contacts, as expected, although apical caps were still visible (Figure 6A-B). Unexpectedly, however, our quantification revealed that the unpolarized localization of aPKC resulted from a loss of apical aPKC, rather than ectopic gain of aPKC at cell-cell contacts (Figure 6B). These data are consistent with the earlier, qualitative description of unpolarized aPKC following PAC-1 depletion, but do not support a model in which loss of *pac-1* causes ectopic aPKC recruitment to cell-cell contacts via ectopic CDC-42 activation.

**Figure 6:**
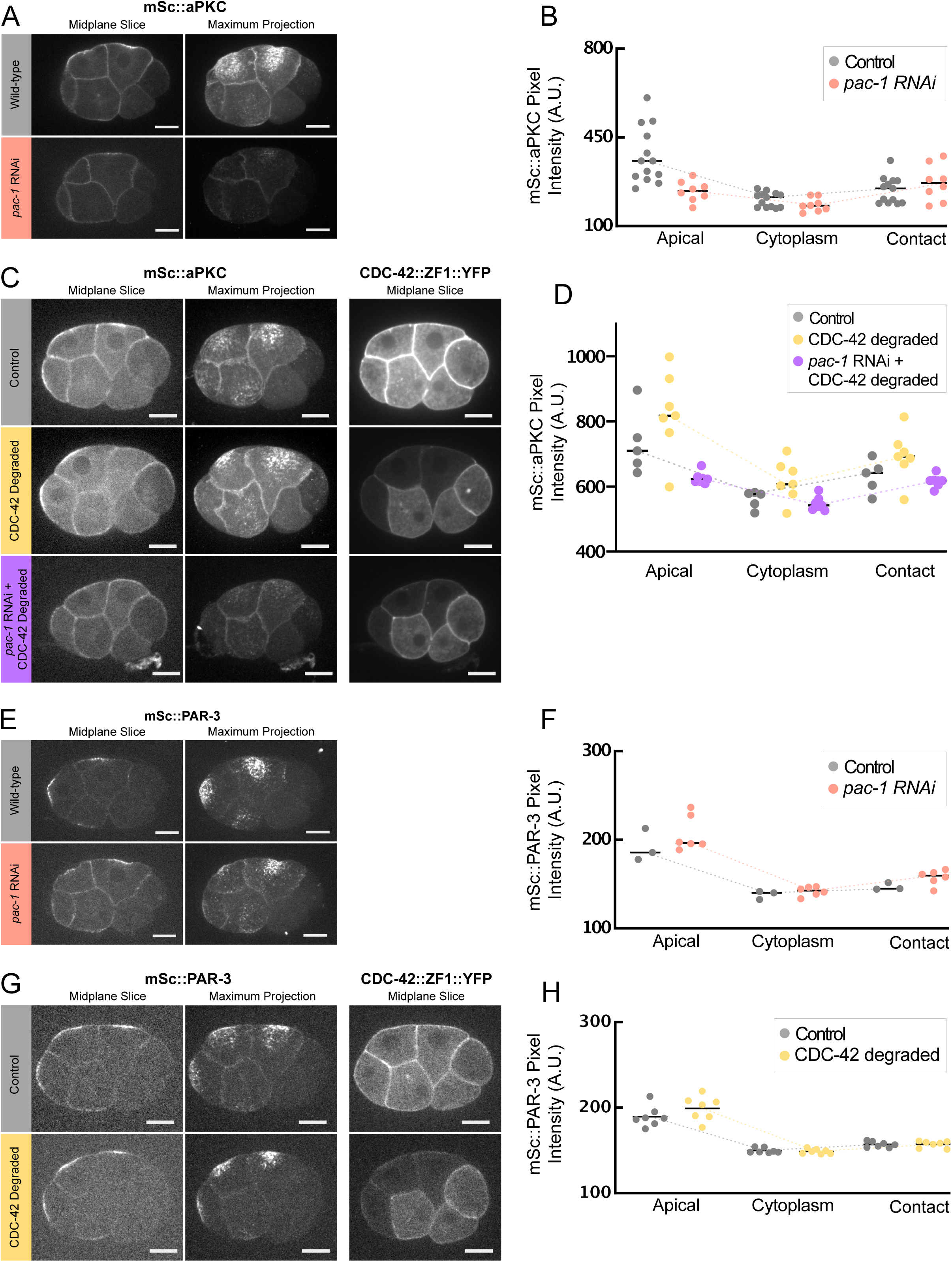
aPKC is regulated by PAC-1 but not through CDC-42. A. mSc::aPKC localization in wild-type and *pac-1* RNAi treated embryos. Midplane and maximum projection shown. Scale bars represent 10 µm. B. Quantification of data in A. Linescans were taken from midplane slices from the apical membrane through the cytoplasm and to the AB cell-cell contact. Apical maximum, cytoplasmic average (of 10 datapoints) and contact maximum plotted. C. mSc::aPKC and CDC-42::ZF1::YFP localization in mSc::aPKC, CDC-42::ZF1::YFP embryos treated with *zif-1* RNAi (no ZF1 degradation; gray), no RNAi (yellow), and *pac-1* RNAi (purple). Scale bars represent 10 µm. D. Quantification of Sc::aPKC fluorescence from linescans of embryos in (C) from apical to contact, as in (B). E. mSc::PAR-3 localization in wild-type and *pac-1* RNAi treated embryos. Midplane and maximum projection shown. Scale bars represent 10 µm. F. Quantification of data in E from linescans from apical, through cytoplasmic to contact. Linescans were taken from midplane slices from the apical membrane through the cytoplasm and to the AB cell-cell contact, as in (B). G. Localization of mSc::PAR-3 and CDC-42::ZF1::GFP. Scale bars represent 10 µm. H. Quantification of data in H from linescans from apical, through cytoplasmic to contact, as in (B). See also: Figure S4

These unexpected results prompted us to re-examine the role of CDC-42 in polarization of aPKC at the 8-cell stage. We reasoned that if the *pac-1* RNAi phenotype was caused by excess CDC-42 activity as previously proposed^22^, then degradation of CDC-42 should at least partly rescue the localization of aPKC in *pac-1(RNAi)* embryos. We therefore repeated *pac-1* RNAi in a strain carrying endogenously tagged YFP::ZF1::CDC-42 to degrade CDC-42 in the AB lineage. CDC-42 levels were reduced ∼75% compared to control embryos treated with *zif-1* RNAi to prevent YFP::ZF1::CDC-42 degradation. In this double depletion experiment, we saw the same phenotype as in *pac-1(RNAi)* alone, with unpolarized aPKC that had equal amounts at caps and at cell-cell contacts (Figure 6C-D, violet).Furthermore, CDC-42 degradation on its own had no effect on aPKC localization to either apical caps or cell-cell contacts (Figure 6C-D, yellow), and CDC-42 itself did not exhibit a visibly polarized localization (Figure 6C, top right panel). Together, these data suggest that CDC-42 does not play an essential role in apical-basal polarity of PAR proteins or apical cap formation at this stage (see Discussion). Because loss of CDC-42 did not rescue the *pac-1* RNAi phenotype, we conclude that PAC-1 must be regulating aPKC via an alternative mechanism that does not involve CDC-42.

We also examined PAR-3 localization and found that it was not measurably affected by depletion of either PAC-1 or CDC-42 (Figure 6E-H). Previously, loss of PAC-1 was reported to cause PAR-3 to be unpolarized on the membrane^22^, but given the dynamic nature of PAR-3 caps during the cell cycle, we suspect that their appearance may have been previously overlooked when examining fixed *pac-1(RNAi)* embryos. Taken together, we conclude that PAR-3 is the most upstream player in apical cap formation (among the proteins we have examined), since it localizes to apical caps independently of both PAC-1 and CDC-42, and partially independently of aPKC. Our data show that PAC-1 controls localization of aPKC to apical caps in a CDC-42-independent manner, perhaps by controlling aPKC recruitment to PAR-3 clusters (see Discussion).

### Apically localized aPKC participates in spindle orientation

The finding that PAC-1 regulates aPKC but not PAR-3 localization at the apical cap provided an opportunity to decouple aPKC localization from PAR-3, allowing us to test whether aPKC and PAR-3 may have different roles in apical cap-dependent spindle orientation. Since aPKC is unpolarized in *pac-1* RNAi (Figure 6A-D), we reasoned that if polarized aPKC was necessary for spindle orientation, there would be a spindle defect in *pac-1* RNAi. Since PAR-3 is unaffected by PAC-1 depletion, if spindle orientation was controlled by PAR-3 independent of aPKC, *pac-1* RNAi embryos would have normal spindle orientation.

We again examined spindle orientation in AB explants and intact embryos by imaging TPXL-1::mSc. The majority of AB explants depleted of PAC-1 formed normal, organized shapes, similar to wild-type, however rare cases of disorganized clusters with 1 or 2 cells without neighbors were still observed (Figure 7A-B, Movie 9). In agreement with this, intact *pac-1(RNAi)* embryos showed signs of spindle angle defects with spindles tilted toward the center of the embryo, although the magnitude of these defects was smaller than in PAR-3::ZF1 or ZF1::aPKC embryos (Figure 7C-E). The severity of the spindle defects in *pac-1(RNAi)* embryos also varied between cells; ABal had defects nearly as severe as those in PAR-3::ZF1 embryos, while spindle angles in ABpr were nearly wild-type (Figure 7D-E, Movie 10) and other cells exhibited intermediate phenotypes (Figures 7D and S5, Movie 10).

**Figure 7:**
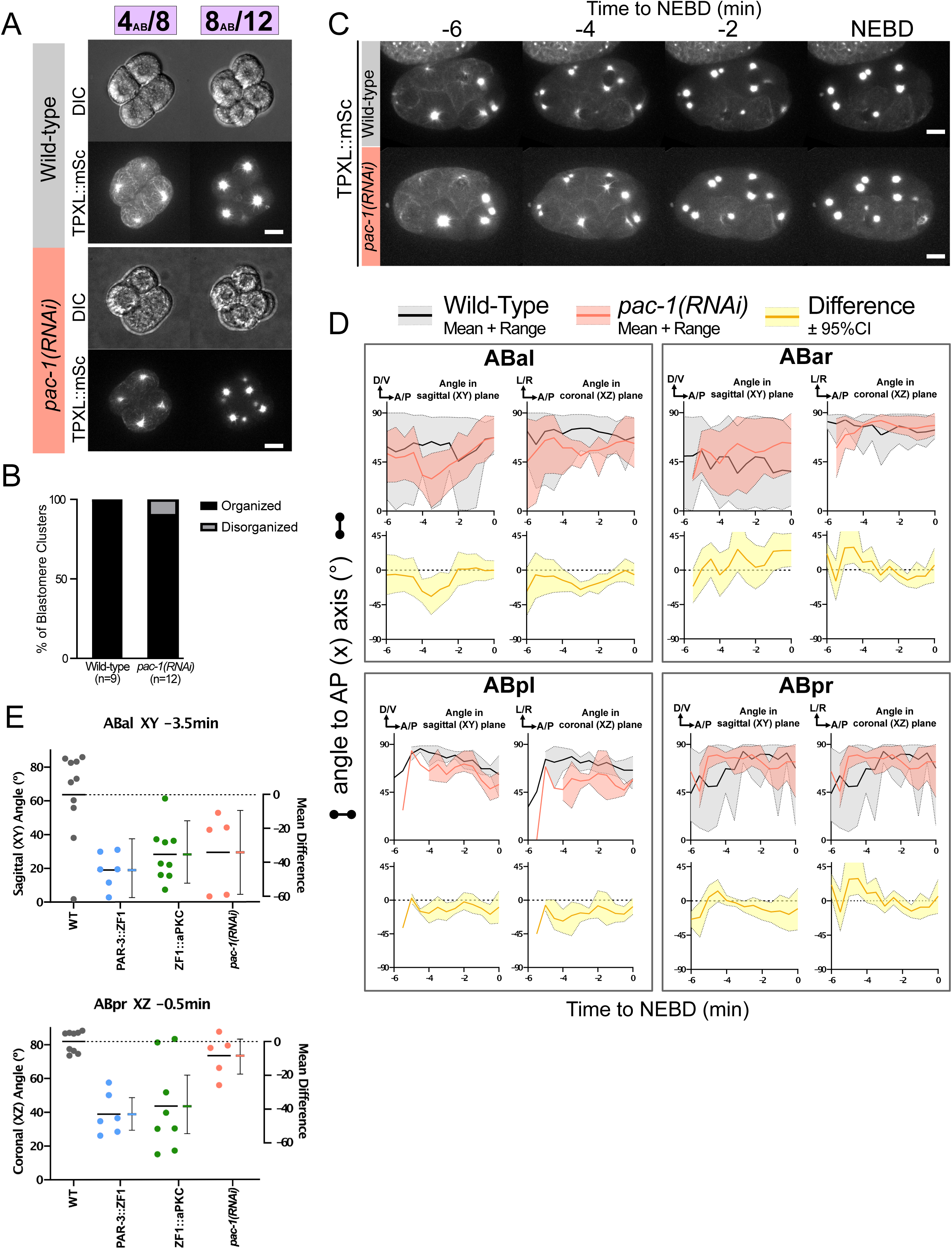
PAC-1 depletion does not cause a spindle orientation defect. A. Representative images of 4_AB_/8 blastomere explants from wild-type and *pac-1* RNAi treated mNG::PAR-3; TPXL-1::mSc embryos. Scale bars represent 10 µm. B. Phenotype scoring of 8_AB_/12 blastomere explants from Figure 7A. Dataset was blinded and scored based on whether the blastomere explant was organized/spherical or disorganized/had cells protruding from the cluster. Wild-type data is the same as in Fig 2 and Fig 5. C. Time-lapse maximum projection montage of centrosomes marked by mSc::TPXL-1 from full-volume imaging in wild-type and *pac-1* RNAi. Time to NEBD (minutes) at top, embryos were imaged every 30 seconds; displayed is every 2 minutes. Scale bars represent 10 µm. D. Quantification of spindle position in ABaL from 3D reconstruction of movies in (C). Centrosomes were annotated and x, y, and z coordinates were used to determine the XY and XZ angles of the spindle relative to the embryo’s AP axis over time, starting from the beginning of centrosome separation until NEBD. Interval = 30sec. E. Comparison of spindle angles between Wild-type, PAR-3::ZF1, ZF1::aPKC and *pac-1(RNAi)* embryos in the indicated cells and at the indicated times. For clarity, we chose to compare the angles and time points at which PAR-3::ZF1 was most different from wild-type. Data points indicate individual measurements, and lines indicate the means. Error bars indicate the mean difference from wild-type and its 95% confidence interval, plotted on the right Y-axis. See Figure S5 for additional cells. See also: Figure S5, Movies 9-10.

Since *pac-1* RNAi did not alter the levels of aPKC at cell-cell contacts, the spindle phenotype of *pac-1* must be due to reduced levels of apical aPKC. We attribute the milder phenotype of *pac-1* (compared to aPKC or PAR-3 depletion) to the fact that apical aPKC was reduced in *pac-1*, rather than eliminated. Together, this indicates that PAR-3 caps alone are not sufficient for normal spindle orientation, but require apical aPKC to be present in order to influence spindle positioning. At the same time, uniformly localized aPKC (in *pac-1(RNAi)*) produced a milder spindle positioning defect than aPKC depletion, suggesting that aPKC contributes to spindle orientation in a manner that does not require aPKC to have a polarized localization.

## Discussion

Inside-outside polarity is a recurring motif in early animal embryonic development, but the different functions of this polarity pattern have not been fully explored. Here, we describe a role for apical PAR-3 caps in oriented, symmetric cell divisions in *C. elegans*. Experiments in mouse embryos have provided a clear connection between apical caps and spindle orientation in determining cell fate^5–7^, but how the polarity proteins work to form apical caps is not clear. Our work here has found an important role for inside-outside polarity in spindle orientation at the 8-cell stage in *C. elegans*, establishing a new context in which apical polarity caps influence development. Previous genetic studies in *C. elegans* outlined some important PAR protein interactions that mediate apicobasal polarity at later stages, where it is important for gastrulation^9, 15, 16, 22–24^. We have shown that the apical domain in 8-cell embryos contains a PAR protein cap that forms in the medial surface during interphase, adding detail and dynamics to the previous characterization of inside-outside polarity. By asking how apical caps form, we found surprising results that caused us to re-evaluate how apicobasal polarity is established in *C. elegans*. We now think of PAR-3 as the central player, whose ability to recruit aPKC apically is regulated in a novel way by PAC-1. Together, these key results provide a new framework for how inside outside polarity can interface with spindle orientation and puts the spotlight on PAR-3 as a key regulator of this process.

We found that when PAR-3 was absent, centrosomes oriented towards cell-cell contacts in both intact embryos and AB explants. From our observations, this phenotype appeared to arise because one centrosome was pulled towards cell-cell contacts in the center of the embryo. We suspect that there is normally force on both centrosomes coming from near the apical cap, which is balanced by forces generated at cell-cell contacts. Consistent with this idea, the apical cap is located directly over the centrosome in each AB blastomere, and fragments of the cap appear to move along with each duplicated centrosome, maintaining a connection as the centrosomes separate (Figure 1D). This model, in which contact-based force on both centrosomes is normally balanced by cap-based force, explains why spindles are more mobile in wild-type than in mutants (Figure 2E, F). An active force balance could enable cells to accurately position spindles in a manner that is flexible and responsive to signaling cues^7^.

Both PAR-3 and aPKC contribute to spindle orientation in the AB cells at the 8-cell stage. The mild spindle orientation defect seen in *pac-1(RNAi)* embryos, where PAR-3 is polarized but aPKC is uniform on the cortex, indicates that polarized PAR-3 alone is not sufficient for proper spindle orientation. Whether the reduction of apical aPKC *pac-1(RNAi)* causes spindle misorientation because aPKC is unpolarized and/or because there is less than some “threshold” amount at the apical surface is an interesting area of further investigation. In one model, apical enrichment of aPKC may be required to achieve an asymmetric distribution of cortical force generators for spindle positioning. There is precedent for localized aPKC regulating spindle pulling forces from the zygote, where aPKC phosphorylates and inhibits NuMA/LIN-5^63^. Alternatively, aPKC may play a more permissive role in controlling spindle orientation rather than providing a spatial landmark, and high apical levels of aPKC are more important than polarized localization *per se.* This second possibility is supported by the observation that degradation of aPKC produced a more severe spindle phenotype than the uniform, unpolarized localization of aPKC seen in *pac-1(RNAi)* embryos. Optogenetic trap experiments to sequester aPKC to certain areas in the cell, as well as experiments to perturb the total levels of aPKC could be starting points to address these ideas.

While aPKC plays a role in spindle orientation, PAR-3 depletion produces the strongest phenotypes in blastomere explants and intact embryos when compared to aPKC depletion and *pac-1(RNAi).* Considering that PAR-3 degradation results in aPKC localizing to the cytoplasm with no obvious membrane accumulation, it is possible that cytoplasmic aPKC results in aberrant/ectopic kinase activity which affects spindles more than aPKC loss. Alternatively, PAR-3 may contribute to spindle orientation via mechanisms that are partially independent from aPKC. Interestingly, in mammalian systems, PAR-3 can interact with microtubule motors, even in the absence of aPKC^64–66^. Since dynein itself does not exhibit an obviously asymmetric localization in 8-cell *C. elegans* blastomeres^38, 67^, we propose that PAR-3 may control dynein motor activity. Alternatively, PAR-3 may act through another motor that does have a polarized localization, or it may indirectly affect motor function, for example by altering microtubule plus-end dynamics at the cortex. If PAR-3 can interact with spindle positioning machinery, the small PAR-3 cap that forms following aPKC degradation could be partially sufficient to orient spindles, which would account for the milder phenotype following aPKC degradation. In the zygote, force imbalances leading to asymmetric spindle orientation have been dissected using laser ablation experiments to sever spindles and observe the recoil/subsequent movements to infer forces on the spindle; similar perturbations could help reveal the mechanistic role of apical caps at later stages^68, 69^.

Although loss of PAR-3 caused spindle orientation defects in both intact embryos and blastomeres, we saw that these defects were corrected prior to cell division in intact embryos, but not in AB explants. We attribute this difference to cell shape; the shapes of cells in intact embryos are influenced by neighbors and confinement within the eggshell, whereas isolated blastomeres are rounder due to lack of confinement. Cell shape is known to influence cell division orientation, with cells generally dividing along the long axis (known as “Hertwig’s rule”). Mechanistic details have been added to this shape-based rule, implicating the ability of longer microtubules to recruit more force generating motors to orient the spindle^59^. This kind of shape-based correction of spindle orientation appears to be pervasive in *C. elegans* embryos; it also happens in 2-cell embryos^60^, in the EMS cell^19, 42^, and in later-stage AB lineage cells^21^. Mitotic correction of initially mis-positioned spindles has also been observed in other systems^61, 62^. Despite the redundancy, our work reveals a form of polarity-dependent spindle positioning that we suspect might be more critical for development in systems where development is more regulative.

In addition to orienting spindles, our results highlight PAR-3’s ability to form apical caps independently of aPKC and CDC-42. Previous experiments in later-stage embryos also showed that PAR-3 could polarize independently of aPKC, but emphasized a putative role for CDC-42 in inside-outside polarity. Specifically, it was proposed that inside-outside polarity is regulated by exclusion of CDC-42, and thus aPARs, at cell-cell contacts, based on the *pac-1* mutant phenotype and the finding that overexpression of CDC-42 variants disrupted polarity^15, 22–24^. We confirmed the key finding that PAC-1 is important for aPKC localization, but its depletion resulted in loss of aPKC apically rather than ectopic accumulation at cell-cell contacts. We also showed that there is a significant pool of aPKC at cell-cell contacts even in wild-type, demonstrating that these cells can polarize without the need to completely exclude aPKC from cell-cell contacts. Together, our results suggest a model in which PAC-1 regulates apical aPKC recruitment through a different small GTPase than CDC-42. This is reasonable since PAC-1 is a rather promiscuous GAP that can also act on other Rho-family small GTPases^70^.

One key difference between our study and previous work is that we focused on the 8-cell stage, whereas Nance and colleagues mainly examined later embryos. Thus, it is possible that the role of CDC-42 is simply different at different stages. Indeed, we know that CDC-42 is required for apical membrane enrichment of myosin during gastrulation^16^, so it clearly has roles in apicobasal polarity in some contexts in the early embryo. The advantage of using 8-cell embryos for mechanistic studies is that we can largely exclude that the phenotypes we observed are a secondary consequence of defects in earlier cell divisions. A caveat is that ZF1-tagged proteins, including CDC-42, are still being actively degraded at the beginning of the cell cycle we are studying, so we cannot rule out that a small amount of remaining CDC-42 is playing a critical role in establishing polarity. Despite this caveat, our data rule out simple models in which each molecule of aPKC/PAR-6 is recruited apically by a molecule of active CDC-42, and they demonstrate that PAC-1’s role in apicobasal polarity cannot be explained solely by its activity towards CDC-42. Our current view of inside-outside polarity at this stage is that PAR-3 instructs aPKC localization at both caps and contacts, however a role for aPKC is still elusive. Presumably, the cap and contact pools of aPKC are tethered to the membrane by different binding partners, thereby modulating different downstream effectors or kinase activity. Our future studies will focus on using single-molecule *ex vivo* biochemistry^32, 71^ to dissect the interactions between PAR proteins during apical cap formation in order to understand how apical caps are formed and how they are regulated by PAC-1.

We were surprised to find that PAR-3 polarization was independent of not only aPKC and CDC-42, but also cytoskeletal inputs including myosin and microtubules (Figure 3). While these results are very different from the zygote, where PAR-3 cannot polarize in the absence of actomyosin flows, they are similar to mouse 8-cell embryos, where it is known that myosin contractility does not directly localize polarity proteins^72^. Microtubules are also dispensable for apical cap formation in the mouse^73, 74^. Our results therefore raise the question of how PAR-3 clusters localize to the center of the apical surface. Cell-cell contacts are known to be important for inside-outside polarity in *C. elegans* embryos based on elegant experiments where two embryos were fused together; apical proteins were excluded from newly formed cell-cell contacts and were localized to the contact free zone^9^. This is an interesting distinction from the mouse, in which single isolated blastomeres can form apical caps^6^. At the outside/apical surface, it is possible that actin-dependent but myosin-independent mechanisms may be at play, as in mice^75, 76^, but further work will be needed to understand how apical cap formation is influenced by cell-cell contacts. Additionally, it is clear that developmental timing plays an important role in both mouse and *C. elegans*, as apical caps first form at the 8-cell stage even though key proteins are present earlier. Stage-specific transcription factors and Rho activation have been implicated in the temporal regulation of cap formation in the mouse embryo^76^, and whether an analogous developmental program governs *C. elegans* 8-cell apical cap formation is an interesting area of investigation. That these multiple inputs work to form the apical cap which then can influence oriented symmetric (described here) and asymmetric (mouse) cell divisions has expanded our view of how apical caps can regulate oriented cell divisions. The ease of working with *C. elegans* embryos, large cell size, and speed over which apical caps form make them an attractive model for understanding the formation and function of apical caps in early animal embryos.

## Acknowledgements

We thank Bruce Bowerman, Jessica Feldman, Ed Munro and Jeremy Nance for sharing *C. elegans* strains; Jacquelyn Wright for assistance with blind scoring of phenotypes; and members of the Dickinson lab for comments on the manuscript. This work was supported by the U.S. National Institutes of Health (R00 GM115964 and R01 GM138443) and by a grant from the Mallinckrodt foundation. NIM was supported by an NIH predoctoral fellowship (F31 HD108006).

DJD is a CPRIT Scholar supported by the Cancer Prevention and Research Institute of Texas (RR170054). Some strains were provided by the Caenorhabditis Genetics Center, which is funded by the NIH Office of Research Infrastructure Programs (P40 OD010440).

## Author Contributions

NJS and DJD conceived of the project and designed the experiments. NIM performed the blastomere isolation experiments, and TS performed the *mrck-1(RNAi)* experiments. NJS performed all other experiments. NJS analyzed the data with assistance from TS and DJD. DJD supervised the project and secured funding. NJS and DJD co-wrote the manuscript, and all authors discussed and contributed to the final version.

## Methods

### Contact for Reagent and Resource Sharing

Requests for resources and further information should be directed and will be fulfilled by the Lead Contact, Daniel J. Dickinson.

### Experimental Model and Subject Details

*C. elegans* strains were fed *E. coli* OP50 on standard NGM growth plates and kept at 20°C unless otherwise noted. Embryos were imaged at a stage before sex determination from hermaphrodites that were not mated to males. A complete list of strains used in this work is included in the Key Resource Table^32, 55, 77–80^.

Genomic editing was performed on *C. elegans* using CRISPR/Cas9 directed homologous recombination using one of three methods:

(1) Injection of plasmids expressing Cas9, guide RNA and repair template^81, 82^ Briefly, for each modification, two plasmids were assembled using PCR amplification and Gibson assembly. These plasmids included expression constructs for Cas9 and the sgRNA sequence corresponding to the locus-of-interest, as well as a homology-directed repair template containing a drug-selectable marker and phenotypic marker. The repair template contained the modification of interest (in this study, fluorescent protein tag) flanked by 500-700bp of homology to the locus of interest. These plasmids, as well as marker plasmids used to exclude animals containing extrachromosomal arrays, were injected into the syncytial gonads of young adult animals. F2 animals were screened for successful Cas9 cutting and homology-directed repair by drug selection and phenotypic marker screening, then the selection markers were removed by Cre-Lox recombination in the F3 generation. Insertion homozygosity was verified by PCR and embryos were imaged to ensure tagged proteins localized and functioned normally.
(2) Injection of Cas9 ribonuclear protein complex and PCR repair template^82^ This method used Cas9 protein, commercially synthesized sgRNA specific to the locus of interest, and a PCR-generated repair template containing the modification of interest (in this study, fluorescent protein tag) with 120bp homology arms corresponding to the locus of interest. The PCR repair template had ssDNA ends made by hybridizing two PCR products of different lengths. The “short” PCR product comprised only the modification of interest (fluorescent protein). The “long” PCR product comprised the modification of interest with 120bp overhangs on the primers that contained homology to the locus of interest. This repair template was purified and mixed with tracrRNA, crRNA, Cas9 protein and a phenotypic marker plasmid. After injection as in (1), F2 insertions were screened by fluorescence of the inserted tag, insertion homozygosity was verified by PCR and embryos were imaged to ensure tagged proteins localized and functioned normally.
(3) Short single-stranded DNA (ssDNA) repair template^83, 84^ This method was used to introduce the myosin temperature-sensitive allele into the strain carrying mNG::PAR-3 and mCherry::Tubulin. Briefly, Cas9 protein, commercially synthesized sgRNA specific to the locus of interest, and commercially-synthesized short ssDNA oligo (ssODN) was used as the repair template. The ssODNcontained the mutation of interest flanked by 30-80nt of homology to the locus of interest, including silent mutations to introduce a new restriction site that enabled screening by restriction fragment length polymorphism (RFLP). In addition, phenotypic markers were added to the injection mix to facilitate screening of edited animals. After screening progeny by RFLP to find putative homozygotes for the edit, the samples were sent for sequencing to verify the point mutation. The new strain was characterized by culturing at the restrictive temperature and determining egg viability, as well as live imaging.

## Method Details

### Confocal microscopy

Embryos were dissected from gravid young adults in a drop of 1x egg buffer (5 mM HEPES pH 7.4, 118 mM NaCl, 40 mM KCl, 3.4 mM MgCl_2_, 3.4 mM CaCl_2_) on a poly-lysine coated 22×22 µM glass coverslip and mounted with agar pads (1.25% agar in 1x egg buffer). Agar pads were made by heating agar on a hotplate, adding one drop from a pasteur pipette to a standard microscope slide, and sandwiching another microscope slide on the agar using two pieces of lab tape as thickness spacers.

Images were collected using one of two instruments. Images in Figures 1C,D; 2E; 3B,C; 5D; and 7C were acquired using a Nikon Ti2 microscope controlled by Micro-Manager software and equipped with a 60x, 1.4 NA oil immersion objective lens; a Visitech iSIM super-resolution confocal scan head; and a Photometrics PrimeBSI or Kinetix22 camera. mNG was excited using 505 nm laser illumination and mCherry, mKate2 and mScarlet-I were excited using 561 nm laser illumination, with appropriate emission filters for detection.mNG was excited using 505 nm laser illumination and mCherry, mKate2 and mScarlet-I were excited using 561 nm laser illumination, with appropriate emission filters for detection. Images in Figures 1B; 2B,C; 3A,D; 4; 5A,B; 6; and 7A were acquired using a Nikon Ti2 microscope controlled by Micro-Manager software and equipped with Nomarski DIC optics for brightfield imaging; a 60x, 1.4 NA oil immersion objective lens; a Crest X-Light V3 spinning disk confocal head; and a Photometrics Prime95B camera. mNG was excited using 488 nm laser illumination and mCherry, mKate2 and mScarlet-I were excited using 555 nm laser illumination, with appropriate emission filters for detection.

### Image processing and display

Images were viewed, processed, analyzed and displayed using the FIJI distribution of ImageJ software. Maximum intensity projections were made where indicated, because apical caps sometimes spanned more than one focal plane. To prepare figures for publication, images were cropped and rotated, and brightness and contrast were adjusted for easy visibility. No other image manipulations were performed.

### Dorsal mounting

Embryos were dissected from gravid adults as above and mounted with 22.8 um beads (Whitehouse scientific, Chester, UK) as spacers. A metal needle was then used to gently slide the cover slip until a 4-cell embryo of interest was rotated with the dorsal side toward the coverslip. Cortical images were acquired as described under “Confocal Microscopy,” above.

### Blastomere isolation

Gravid adults were dissected in 1x egg buffer. Immediately after dissecting, 10% sodium hypochlorite (Sigma 239305) was added for 2-3 min to remove the vitelline layer. Embryos were then washed in egg buffer followed by Shelton’s growth media (SGM) supplemented with 35% fetal bovine serum. Washes were carried out by using a mouth pipet to transfer embryos from one drop of solution to another in a depression slide. To remove the chitin layer, embryos were transferred into a solution of 10 units/mL chitinase (SIGMA-C6137) mixed with 10 mg/mL chymotrypsin (Sigma C4129) for 7mins. De-shelled embryos were then washed 3x in SGM. The anterior blastomere (AB) was dissociated from each 2-cell stage embryo by mouth pipetting using a glass needle. Needles were made by pulling glass capillary tubes (World precision instrument IB100F-4) in a needle puller (Sutter instrument P-97) and were trimmed under a micro forge microscope (Narishige MF-9) to an inside diameter of 250-300 µm. The isolated AB was mounted in SGM containing polystyrene microspheres (Cospheric PSMS-1.07 14-20um) for confocal imaging.

### Full volume imaging, image processing for 3D rendering, and quantification of spindle angles

For full volume imaging of embryos to measure spindle angle, embryos were mounted with coverslips as above, and 0.2um step-size Z-slices were acquired through the full embryo volume, between 80-100 Z-slices. Using FIJI, images were rotated to make EMS spindle at cytokinesis 0° relative to the AP axis. Background subtraction was performed using the “Process>Subtract background” command in FIJI, with the default 50 pixel rolling ball radius. To create a binary image, images were thresholded using “Image>Adjust>Auto threshold” with the “Max Entropy” filter, with “white on black” and “stack” selected. Thresholded images were opened in 3D viewer (“Plugins>3D Viewer”). To get x, y, and z positions of each centrosome over time, the “multipoint” tool was used to mark each centrosome.

The XY angle *θ* between the anterior-posterior axis and the line connecting the two centrosomes was calculated from the x and y coordinates of each centrosome from Fiji 3D viewer annotation:

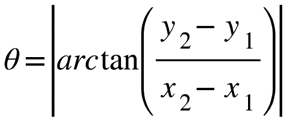

The XZ angle *φ* between the anterior-posterior axis and the line connecting the two centrosomes was calculated from the x and y coordinates of each centrosome from Fiji 3D viewer annotation:

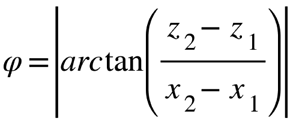

The 3D centrosome distance was calculated using the distance formula in three dimensions where *CD* is the distance between the two centrosomes which each have 3-dimensional coordinates *C*(x_1_, y_1_, z_1_) and *D*(x_2_, y_2_, z_2_)

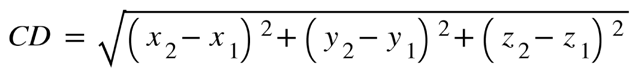

To assess the magnitude of each perturbation’s effect on spindle angles, we calculated the effect size (difference between means of wild-type and each mutant as a function of time) and its 95% confidence interval using the DABEST package for MATLAB ^85, 86^.

### Drug treatment

L4 and young adult worms were fed *perm-1* RNAi^87^ for 17-20 hours. To test for permeability, embryos were dissected from gravid adults in Shelton’s Growth Media with 35% FBS (SGM) and transferred to a drop of bleach solution to monitor for cytoplasmic clearing. Fresh adult worms were dissected into SGM + 0.5% DMSO on a 24×60mm polylysine-coated coverslip and mounted with 22.8um glass beads with the slide and coverslip arranged perpendicular to one another. The two sides where the glass slide overhangs the coverslip were sealed. The sides with the coverslip overhanging the slide were left unsealed to serve as an inlet/outlet of a makeshift flow cell. To facilitate wicking, 3-5 µL of SGM was added to the coverslip at the inlet and outlet just prior to imaging. Embryos were imaged using the spinning disk confocal system described above, and nocodazole was added just after the 4-8 cell AB division. To add nocodazole, 3-5 µL of SGM + 50 µM nocodazole (dissolved in DMSO) was pipetted onto one side of the open coverslip and wicked through using a Kimwipe placed on the other open side.

### RNA interference

RNA interference (RNAi) was performed by injection (Figure 6A) or by feeding (all other experiments). For *pac-1,* where a mixture of injection and feeding experiments, we confirmed that both methods affected aPKC localization equivalently. For injection, 0.5-1.5 kb of the target gene was amplified from N2 cDNA with primers including T7 promoters. ssRNA was transcribed from this PCR product using the Promega T7 RiboMAX Express kit (Cat. No. P1700), purified, and annealed to form dsRNA. dsRNA was injected into the body of young adult worms at 1µg/µL and worms were transferred to standard OP50 NGM plates for 24-28 hours before dissection and embryo imaging.

For feeding, bacteria clones expressing dsRNA from the *C. elegans* RNAi feeding library (ref) were grown and sequence verified, then 5 mL of culture was grown for 8 hours, concentrated into 1mL, and 50 uL of this suspension was plated onto NGM plates containing carbenicillin and IPTG to induce dsRNA expression. After bacteria growth and dsRNA induction for at least 4 hours, L4/young adult worms were transferred onto the feeding plates. *perm-1* dsRNA was fed for 17-20 hours to reduce pleiotropic effects seen with longer feeding times. *zif-1* and *pac-1* dsRNAs were fed for 24-28 hours.

### Myosin ts experiments

Worms were cultured at 15°C. To maintain reagents at a similar temperature at the bench, a small styrofoam cooler outfitted with a thermometer was filled with ice, and the ice was topped with plastic pipette tip boxes to serve as insulation. Agar pads, depression slides (for dissection) and 1x egg buffer were chilled in the cooler on the plastic (to avoid direct contact with the ice) where the ambient temperature was measured to be between 10-15°C. Working quickly, one polylysine coverslip was placed on top of a chilled depression slide, 5 µL chilled egg buffer was added, and worms were dissected. The coverslip was mounted with a pre-chilled agar pad, then transferred to a stage-top incubator installed on the spinning disk confocal microscope (described above), which was pre-heated to 25°C using an objective heater collar. Images were collected as quickly as possible after temperature shifting.

### Quantification of apical, cytoplasmic and cell-cell contact fluorescence intensities

Apical, cytoplasmic, and contact fluorescence intensity was quantified in FIJI. A 25 pixel-wide segmented line was drawn from outside the embryo, through the apical surface perpendicularly, through the cytoplasm around the nucleus, and through the nearest AB cell-cell contact. The pixel intensity across the line distance was plotted using “Analyze>Plot Profile.” The apical maximum was determined by finding the midpoint of the dataset and determining the maximum value in the first half of the dataset. The contact maximum was found by finding the maximum value in the second half of the dataset. The cytoplasmic value was calculated by averaging 10 points in the middle of the dataset.

## Key Resource Table

|  |  |  |  |  |
| --- | --- | --- | --- | --- |
| Chemicals |  |  |  |  |
| Reagent | Source | Identifier |  |  |
| Nocodazole | Enzo | Catalog number: BML-T101-0010 |  |  |
| Experimental Models: *C. elegans* strains |  |  |  |  |
| Strain name | Description | Genotype | Source | Construction Method |
| N2 | Wild-type |  | Caenorhabditis Genetics Center |  |
| EU1627 | mCh::Tubulin | *unc-119(ed3) III; orEx22[tba-1p::mCherry::tba-1(genomic)+unc119(+)]* | O'Rourke et al. 2010 |  |
| LP242 | mNG::PAR-3 | *par-3(cp54[mNeonGreen::3xFlag::par-3]) III* | Dickinson et al. 2017 |  |
| LP427 | mNG::PAR-3; mCh::Tubulin | *par-3(cp54[mNeonGreen::3xFlag::par-3]) III; orEx22[tba-1p::mCherry::tba-1(genomic)+unc119(+)]* | This work | Cross between LP242 and EU1627 |
| UTX134 | TPXL-1::mSc | *tpxl-1(djd41[tpxl-1::mSc]) I* | This work | Cas9 RNP injection into N2 with PCR repair template |
| UTX141 | TPXL-1::mSc; mNG::PAR-3 | *tpxl-1(djd41[tpxl-1::mSc]) I; par-3(cp54[mNeonGreen::3xFlag::par-3]) III* | This work | Cross between LP242 and UTX134 |
| JLF384 | par-3::zf:gfp:3xflag exon 6a | *par-3(wow59[par-3::zf:gfp:3xflag exon 6a]) zif-1(gk117)* | Pickett et al. 2022 |  |
| UTX175 | TPXL-1::mSc; PAR-3::ZF1::GFP exon 6a | *tpxl-1(djd41[tpxl-1::mSc]) I; par-3(wow59[par-3::zf:gfp:3xflag exon 6a]) III* | This work | Cross between JLF384 and UTX134 |
| LP636 | mNG::PAR-3; NMY-2::mKate2 | *nmy-2(cp52[nmy-2::mkate2 + LoxP unc-119(+)] LoxP) I; par-3(cp54[mNeonGreen::3xFlag::par-3]) III* | Dickinson et al. 2017 |  |
| UTX224 | PAR-3::mNG; mCh::Tubulin; NMY-2(ts) | *nmy-2(djd57[nmy-2::ts]) I.; par-3(cp54[mNeonGreen::3xFlag::par-3]) III; Ppie-1::mCherry::TBA-1* | This work | Cas9 RNP injection into LP427 with ssODN repair template |
| JLF504 | par-1::zf:gfp:3xflag exon 11a; zif-1(gk117) | *par-1(wow87[par-1::zf:gfp:3xflag exon 11a]); zif-1(gk117)* | Pickett et al. 2022 |  |
| UTX58 | mSc::aPKC | *pkc-3(djd15[mSc::Myc::aPKC]) II* | This work | Cas9 RNP injection into N2 with PCR repair template |
| UTX129 | mSc::aPKC; PAR-1::ZF1::GFP | *pkc-3(djd15[mSc::Myc::aPKC]) II; par-1(wow87[par-1::zf:gfp:3xflag exon 11a]) V.* | This work | Cross between JLF504 and UTX58 |
| JLF480 | ZF1::GFP::aPKC | *pkc-3(wow85[zf:gfp:3xflag:pkc-3]); zif-1(gk117)* | Sallee et al. 2021 |  |
| UTX113 | mSc::PAR-3 | *par-3(djd33[mSc-I-GLO::Myc::par-3]) III* | Chang & Dickinson 2022 |  |
| UTX171 | ZF1::GFP::aPKC; Sc::PAR-3 | *pkc-3(wow85[zf:gfp:3xflag:pkc-3]) II; par-3(djd33[mSc-I-GLO::Myc::par-3]) III* | This work | Cross between JLF480 and UTX113 |
| UTX178 | TPXL-1::mSc; ZF1::GFP::aPKC | *tpxl-1(djd41[tpxl-1::mSc]) I; pkc-3(wow85[zf:gfp:3xflag:pkc-3]) II.* | This work | Cross between JLF480 and UTX134 |
| FT1896 | ZF1::YFP::CDC-42 | *cdc-42(xn65[zf1::yfp::cdc-42 + LoxP::unc-119::LoxP]) II* | Abrams & Nance 2021 |  |
| UTX170 | mSc::PAR-3; ZF1::YFP::CDC-42 | *cdc-42(xn65[zf1::yfp::cdc-42 + LoxP::unc-119::LoxP]) II; par-3(djd33[mSc-I-GLO::Myc::par-3]) III* | This work | Cross between FT1896 and UTX113 |
| UTX172 | mSc::aPKC; ZF1::YFP::CDC-42 | *pkc-3(djd15[mSc::Myc::aPKC]), cdc-42(xn65[zf1::yfp::cdc-42 + LoxP::unc-119::LoxP]) II.* | This work | Cross between FT1896 and UTX58 |
| UTX131 | mSC::PAR-3; PAR-1::ZF1::GFP | *par-3(djd33[mSc-I-GLO::Myc::par-3]) III; par-1(wow87[par-1::zf:gfp:3xflag exon 11a]) V* | This work | Cross between JLF504 and UTX113 |
| Software and Algorithms |  |  |  |  |
| Resource name | Source | Reference |  |  |
| DABEST for MATLAB | https://github.com/ACCLAB/DABEST-Matlab | Ho et al. 2019 |  |  |

---

**Figure S1.**
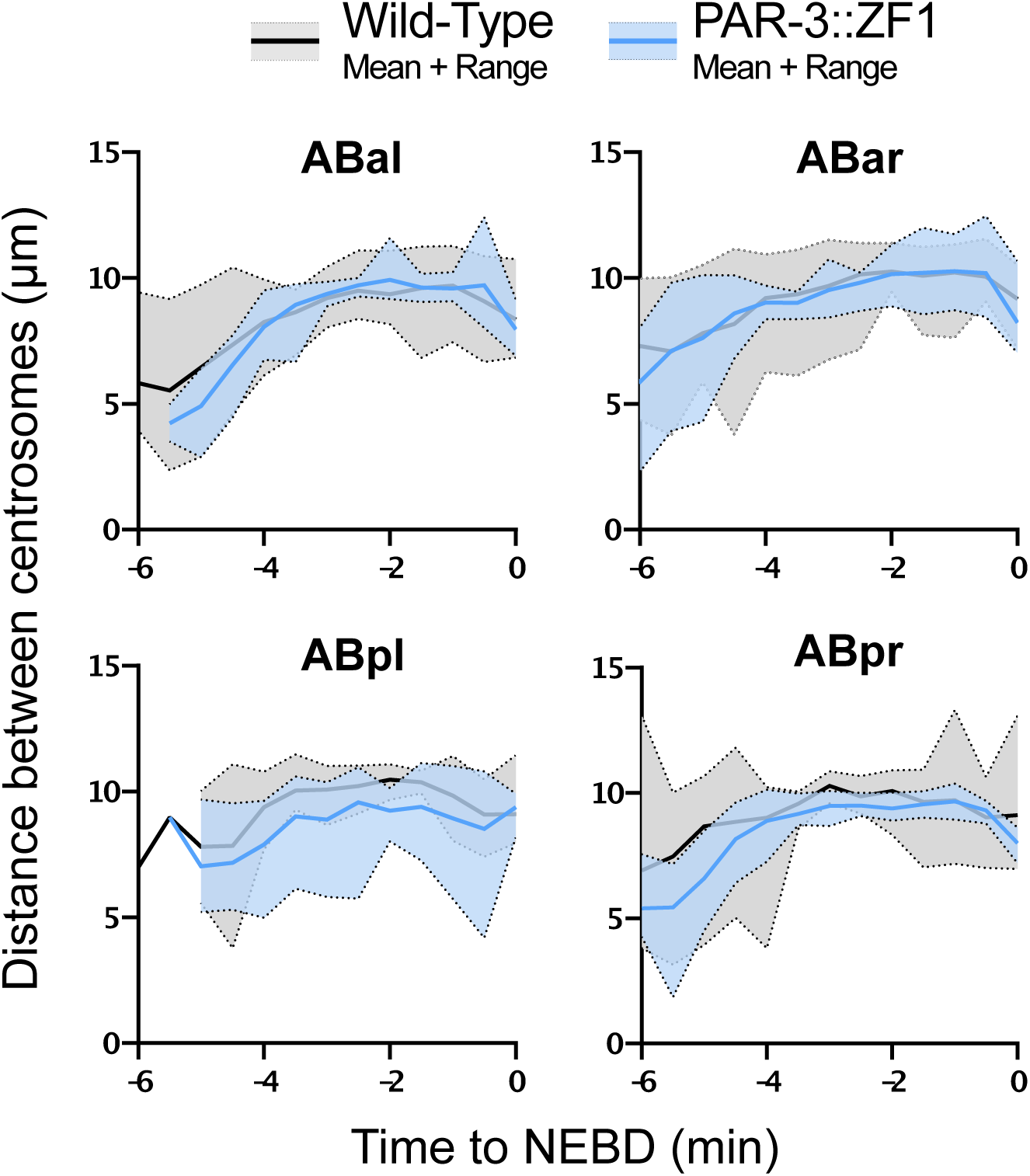
related to Figure 2: Quantification of centrosome separation in PAR-3::ZF1. Full-volume images were displayed in 3D, and centrosomes were annotated manually. X, Y, and Z coordinates were used to determine the 3D distance between centrosomes over time, starting from the beginning of centrosome separation until NEBD. Time is shown relative to NEBD.

**Figure S2.**
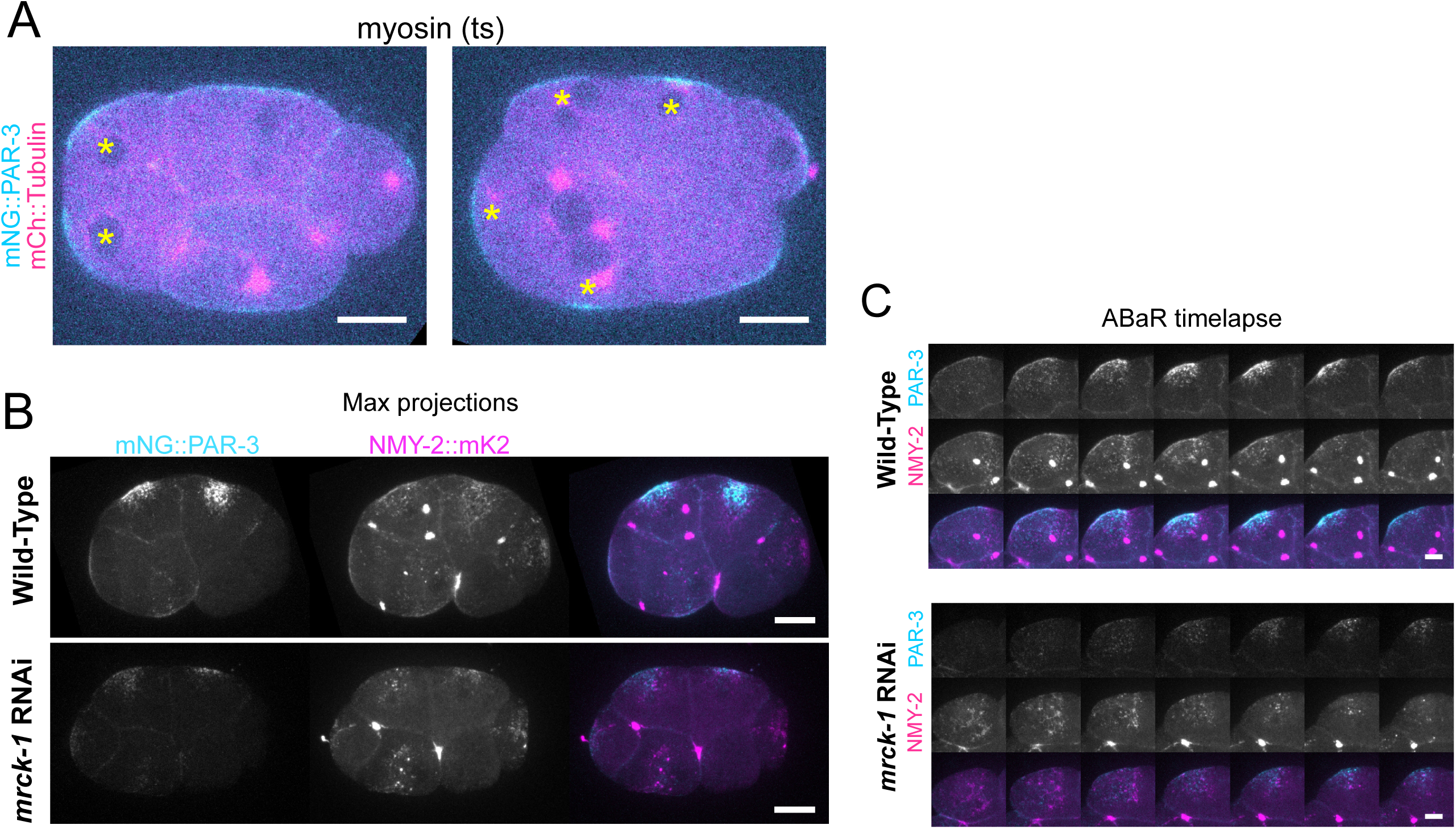
Related to Figure 3: *mrck-1(RNAi)* disrupts apical myosin but not apical PAR-3 caps. A. Images from myosin (ts) (4C) that fail cytokinesis and form multinucleate cells. We saw PAR-3 accumulations near these extra nuclei/centrosomes even in cases where there was no membrane in between (as in left image), suggesting some connection between nuclei/centrosomes/MTOC and PAR-3 accumulation. Scale bars represent 10 µm. B. mNG::PAR-3 and NMY-2::mK2 localization in wildtype and *mrck-1* RNAi treated embryos. Max projections are shown. Scale bars represent 10 µm. C. Montage of ABar in wild-type and *mrck-1* RNAi embryos. Scale bars represent 5 µm.

**Figure S3.**
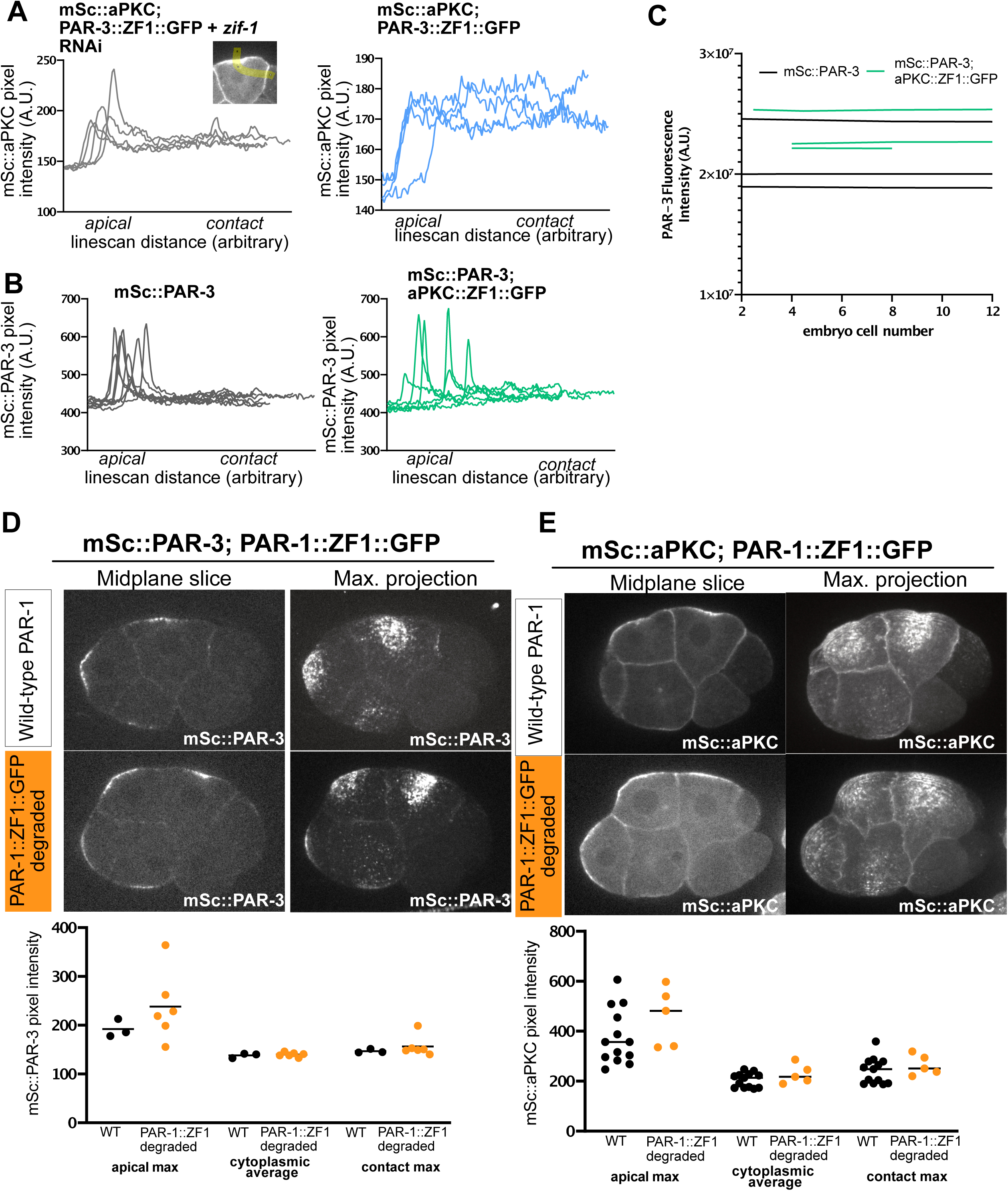
Related to Figure 4: Apical cap formation is independent of PAR-1. A. Line scans used for quantification in Figure 4C (mSc::aPKC in PAR-3::ZF1::GFP with or without *zif-1(RNAi)* to prevent PAR-3 degradation). B. Line scans used for quantification in Figure 4E (mSc::PAR-3 in wild-type vs. aPKC::ZF1::GFP) C. mSc::PAR-3 fluorescence intensity over time in aPKC::ZF1::GFP, showing that PAR-3 levels are not reduced when aPKC is degraded. D. Top: Localization of mSc::PAR-3 in a wild-type or PAR-1::ZF1 background. Bottom: Quantification as illustrated in Figure 4B. E. Top: Localization of mSc::aPKC in a wild-type or PAR-1::ZF1 background. Bottom: Quantification as illustrated in Figure 4B.

**Figure S4.**
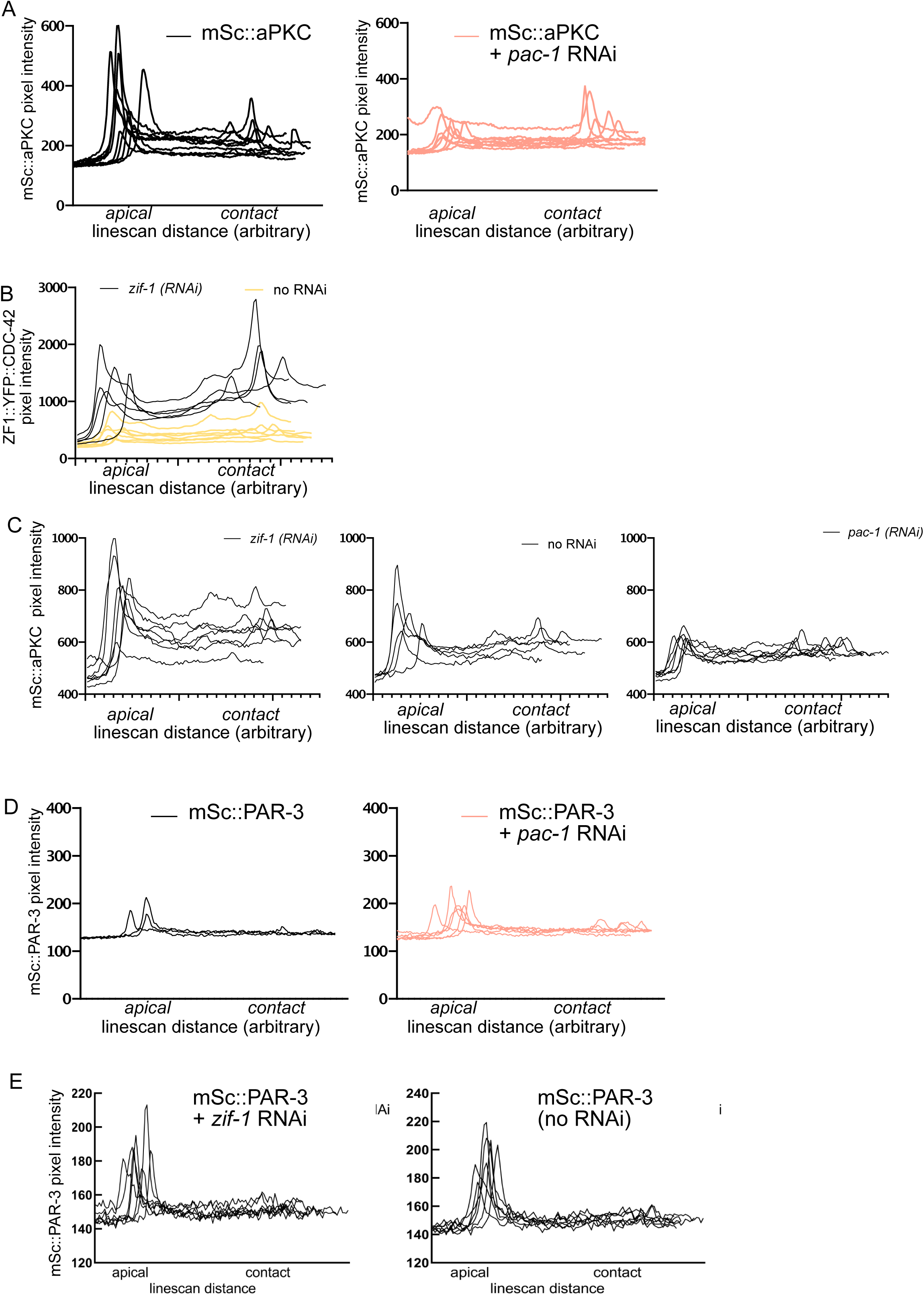
Related to Figure 6: Quantification of protein localizations for Figure 6. A. Line scans used for quantification in Figure 6B (mSc::aPKC in wild-type vs. *pac-1(RNAi)*). B. Line scans showing the extent of CDC-42 depletion in Sc::aPKC; CDC-42::ZF1::YFP. C. Line scans used for quantification in FIgure 6D (mSc::aPKC in CDC-42::ZF1::YFP, with or without *pac-1(RNAi)* and with or without *zif-1(RNAi)* to prevent CDC-42 degradation). D. Line scans used for quantification in Figure 6F (mSc::PAR-3 with or without *pac-1(RNAi)*). E. Line scans used for quantification in Figure 6H (mSc::PAR-3 in CDC-42::ZF1::YFP, with or without *zif-1(RNAi)* to prevent CDC-42 degradation).

**Figure S5.**
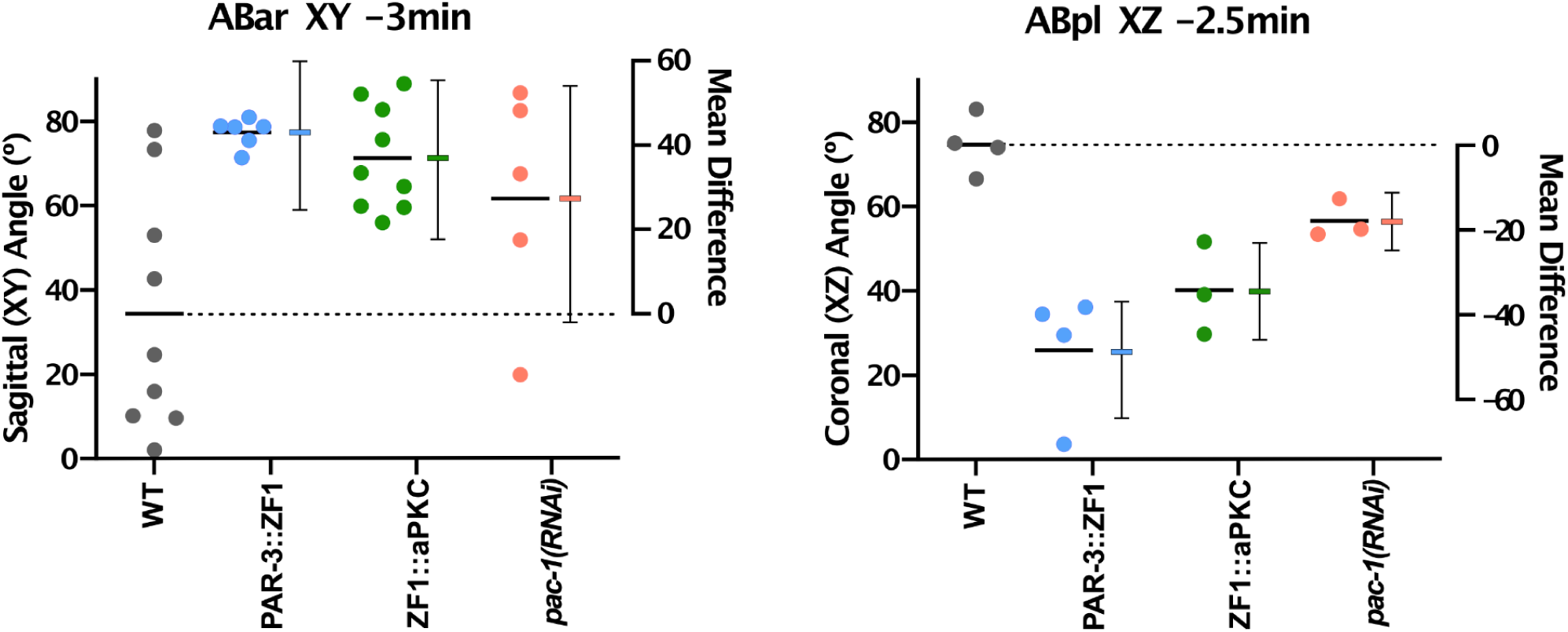
related to Figure 7: Additional comparisons between different genotypes. Comparison of spindle angles between Wild-type, PAR-3::ZF1, ZF1::aPKC and *pac-1(RNAi)* embryos in the indicated cells and at the indicated times. For clarity, we chose to compare the angles and time points at which PAR-3::ZF1 was most different from wild-type. Data points indicate individual measurements, and lines indicate the means. Error bars indicate the mean difference from wild-type and its 95% confidence interval, plotted on the right Y-axis.

## Movies

Movie 1, Related to Figure 1B:

Time-lapse movie of mNG::PAR-3 and mCh::Tubulin tagged embryo viewed from the left, ABal and ABpl are visible along the top & left of the embryo. Interval = 30 seconds.

Movie 2, Related to Figure 1D:

Time-lapse movie of embryo expressing PAR-3::mNG and mCh::Tubulin, mounted en-face dorsally with the ABal blastomere in view. Movie starts just after ABxx cells are born. Interval =1.5 seconds.

Movie 3, Related to Figure 2C:

Time-lapse movie of blastomere explants from wild-type (mNG::PAR-3; TPXL-1::mSc) imaged from 4_AB_/8 to 8_AB_/12 stage. DIC, mNG::PAR-3 and TPXL-1::mSc dataset are shown. Interval = 2 minutes.

Movie 4, Related to Figure 2C:

Time-lapse live movies of blastomere explants from PAR-3::ZF1::GFP (PAR-3::ZF1::GFP; TPXL-1::mSc) imaged from 4_AB_/8 to 8_AB_/12 stage. DIC, PAR-3::ZF1::GFP and TPXL-1::mSc are shown. Interval = 2 minutes.

Movie 5, Related to Figure 2D:

TPXL-1::mSc maximum projection images from time-lapse live full-volume imaging of wild-type (mNG::PAR-3; TPXL-1::mSc) embryos. Right side is displayed, three ABxx cells are visible along the top and left of the embryo. Interval = 30 seconds.

Movie 6, Related to Figure 2D:

TPXL-1::mSc maximum projection images from time-lapse live full-volume imaging of

PAR-3::ZF1::GFP (PAR-3::ZF1::GFP; TPXL-1::mSc) embryos. Right side is displayed, three ABxx cells are visible along the top and left of the embryo. Interval = 30 seconds.

Movie 7, Related to Figure 5B:

Time-lapse movie of blastomere explants from ZF1::GFP::aPKC (ZF1::GFP::aPKC; TPXL-1::mSc) embryo from the 4_AB_/8 to the 8_AB_/12 stage. Interval = 2 minutes.

Movie 8, Related to Figure 5D:

TPXL-1::mSc maximum projection time-lapse movie from full-volume imaging of ZF1::GFP::aPKC (ZF1::GFP::aPKC; TPXL-1::mSc) embryos. Right side is displayed, three ABxx cells are visible along the top and left of the embryo. Interval = 30 seconds.

Movie 9, Related to Figure 7B:

Time-lapse movie of blastomere explants from *pac-1(RNAi)* (mNG::PAR-3; TPXL-1::mSc) embryos from the 4_AB_/8 to the 8_AB_/12 stage. Interval = 2 minutes.

Movie 10, Related to Figure 7D:

TPXL-1::mSc maximum projection time-lapse movie from full-volume imaging of*pac-1(RNAi)* (mNG::PAR-3; TPXL-1::mSc) embryos. Right side is displayed, three ABxx cells are visible along the top and left of the embryo. Interval = 30 seconds.

